# Biophysical network modeling of temporal and stereotyped sequence propagation of neural activity in the premotor nucleus HVC

**DOI:** 10.1101/2024.11.13.623378

**Authors:** Zeina Bou Diab, Marc Chammas, Arij Daou

## Abstract

Stereotyped neural sequences are often exhibited in the brain, yet the neurophysiological mechanisms underlying their generation are not fully understood. Birdsong is a prominent model to study such behavior particularly because juvenile songbirds progressively learn from their tutors and by adulthood are able to sing stereotyped song patterns. The songbird premotor nucleus HVC coordinate motor and auditory activity responsible for learned vocalizations. The HVC comprises three neural populations that has distinct *in vitro* and *in vivo* electrophysiological responses. Typically, models that explain HVC’s network either rely on intrinsic HVC circuitry to propagate sequential activity, rely on extrinsic feedback to advance the sequence or rely on both. Here, we developed a physiologically realistic neural network model incorporating the three classes of HVC neurons based on the ion channels and the synaptic currents that had been pharmacologically identified. Our model is based on a feedforward chain of microcircuits that encode for the different sub-syllabic segments (SSSs) and that interact with each other through structured feedback inhibition. The network reproduced the *in vivo* activity patterns of each class of HVC neurons, and unveiled key intrinsic and synaptic mechanisms that govern the sequential propagation of neural activity by highlighting important roles for the T-type Ca^2+^ current, Ca^2+^-dependent K^+^ current, A-type K^+^ current, hyperpolarization activated inward current, as well as excitatory and inhibitory synaptic currents. The result is a biophysically realistic model that suggests an improved characterization of the HVC network responsible for song production in the songbird.

**Significance Statement:** Learned motor sequences acquired through repetitive practice undergo stabilization and are integrated into cortical motor circuits through a process of consolidation. The process of mastering a complicated motor sequence requires thorough motor exploration to achieve closer alignment with the desired outcome, yet the neural mechanisms underlying sequence generation remain largely unexplored. In this study, we investigate the neural circuitry in a premotor region of the songbird brain (known as HVC) that is well-known for generating extremely precise learned sequences. We develop a biophysically realistic network model that incorporate pharmacologically identified intrinsic and synaptic currents for the three different classes of HVC neurons. The network highlights fundamental intrinsic and synaptic mechanisms regulating the sequential propagation of neural activity.

## Introduction

Learned temporal sequences are expressed in many brain regions and play a critical role in temporal information encoding, such as navigation (Skaggs et al., 1996), the timing of motor actions (Wang et al., 2018), time generation (MacDonald et al., 2011; Pastalkova et al., 2008), decision making (Harvey et al., 2012; Schmitt et al., 2017) and skilled movement (Peters et al., 2014). Zebra finches sing remarkably stereotyped songs rendering them as an excellent model for studying the mechanisms underlying neural sequences (Hahnloser et al., 2002). The anatomical basis for song production and learning is a highly developed neural network known as the song system (Fig. 1A), with nucleus HVC exhibiting a rhythmic pattern-generating role encoding for the syllable order and the overall temporal structure of the birdsong (Fee & Goldberg, 2011; Fee et al., 2004; Fee & Scharff, 2010; Mooney, 2009; Yu & Margoliash, 1996).

**Figure 1.**
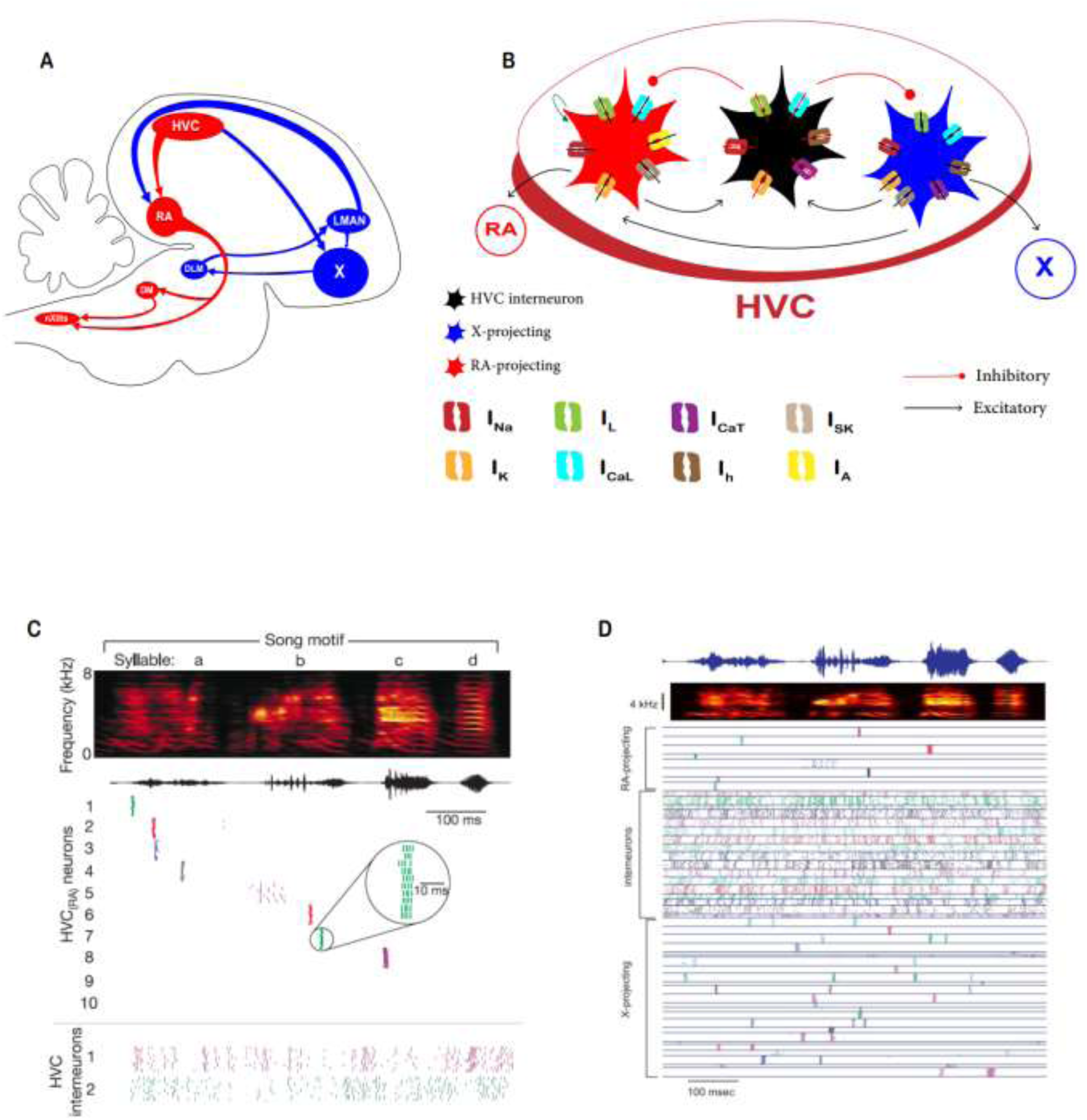
Song system overview and general intrinsic and network properties of HVC neurons in vivo and in vitro. **A**. Schematic diagram showing a sagittal view of the male zebra finch song system. The vocal motor pathway (VMP, red color) contains circuits that directly pattern song output. The anterior forebrain loop (AFP, blue color) pathway contains circuits that are important for song learning and plasticity. **B**. HVC includes multiple classes of neurons; HVC_X_ neurons that project to area X (blue), HVC_RA_ neurons that project to nucleus RA (red) and HVC interneurons (HVC_INT_, black). HVC_X_ and HVC_RA_ excite HVC_INT_ via AMPA and NMDA synapses (green arrow), while HVC_INT_ neurons inhibit both classes of projecting neurons via GABA synapses (brown arrows with circle heads). Each class of HVC neurons is characterized with its own family of ionic currents (Daou et al., 2013). **C**. HVC_RA_ neurons exhibit a very sparse activity during singing eliciting a single 4-6 ms burst at a single and exact moment in time during each rendition of the song. On the contrary, HVC interneurons burst densely throughout the song (Adopted from (Hahnloser et al., 2002)). **D**. Similar to HVC_RA_, HVC_X_ neurons generate 1-4 bursts that are time-locked and highly stereotyped from one rendition of the song to another (Adopted from (Kozhevnikov & Fee, 2007)).

There are three main neuronal populations in the HVC exhibiting different functional, cellular and pharmacological properties (Daou et al., 2013; Kubota & Taniguchi, 1998; Mooney, 2000; Mooney & Prather, 2005): neurons that project to the robust nucleus of arcopallium (RA; HVC_RA_), neurons that project to Area X (HVC_X_), and interneurons (HVC_INT_, Fig. 1B). HVC_Av_ neurons (projecting to nucleus Avalanche) exists and play a role in song learning, yet their intrinsic and synaptic properties remains unknown (Roberts et al., 2017). Intracellular recordings showed that HVC_RA_, HVC_X_, and HVC_INT_ neurons have distinctive *in vivo* and *in vitro* electrophysiological properties (Daou et al., 2013; Dutar et al., 1998; Kubota & Taniguchi, 1998; Mooney et al., 2001; Shea et al., 2010), which are orchestrated via a family of ion channels (Daou et al., 2013).

During singing, HVC_RA_ neurons produce a single burst of spikes (∼10 ms) that are tightly locked to the song (Fig. 1C, (Hahnloser et al., 2002; Kozhevnikov & Fee, 2007). HVC_X_ neurons in their turn elicit 1-4 bursts that are also time-locked to vocalizations (Fig. 1D, (Kozhevnikov & Fee, 2007), while HVC_INT_ neurons exhibit tonic activation (Fig. 1C), with bursting and suppression at different locations throughout the song (Amador et al., 2013; Cannon et al., 2015; Hahnloser et al., 2002; Kosche et al., 2015; Kozhevnikov & Fee, 2007; Long et al., 2010; Markowitz et al., 2015).

Several models of how sequence is generated within HVC have been proposed (Cannon et al., 2015; Drew & Abbott, 2003; Egger et al., 2020; Elmaleh et al., 2021; Galvis et al., 2018; Gibb et al., 2009a, 2009b; Hamaguchi et al., 2016; Jin, 2009; Long & Fee, 2008; Markowitz et al., 2015). These models either rely on intrinsic HVC circuitry to propagate sequential activity, rely on extrinsic feedback to advance the sequence or rely on both. The proposed models do not capture the complex details of spike morphology, do not include the right ionic currents, do not incorporate all classes of HVC neurons, or do not generate realistic firing patterns as seen *in vivo*. In this work, we ask a simple but powerful question: Can the biologically realistic intrinsic properties of HVC neurons and the local synaptic connections among them produce the precise sequence propagation seen in birdsong? We are particularly interested in understanding what core biophysical ingredients (ionic and synaptic currents) are truly necessary to generate sparse sequences while preserving each HVC neuron activity in vivo. How do features like bursting behavior of these neurons or patterns of inhibition help shape the flow of activity throughout network?” To address these questions, we developed a physiologically realistic network model incorporating the three classes of HVC neurons based on the ion channels and the synaptic currents that had been pharmacologically identified (Daou et al., 2013; Kosche et al., 2015; Mooney & Prather, 2005). Our model is based on a feedforward chain of microcircuits that encode for the different sub-syllabic segments (SSSs) and that interact with each other through structured feedback inhibition. The network developed unveiled key intrinsic and synaptic mechanisms that govern the sequential propagation of neural activity by highlighting important roles for the T-type Ca^2+^ current, Ca^2+^-dependent K^+^ current, A-type K^+^ current, hyperpolarization activated inward current, as well as excitatory and inhibitory synaptic currents. Our model provides a new way of thinking about sequence generation during birdsong vocalizations and in network architectures more generally.

## Methods

Single-compartment conductance-based Hodgkin-Huxley-type (HH) biophysical models of cells from the HVC were developed and connected together via biologically realistic synaptic currents. Simulations of these model neurons and of the model network composed of synaptically coupled HVC_RA_, HVC_X_, and HVC_INT_ neurons were performed using the ode45 numerical integrator in MATLAB (<u>MathWorks</u>). Source codes for each network will be made available online at our lab’s website as well as on ModelDB.

HVC model cells that are used to connect the networks exhibited ionic and synaptic currents that had been shown to be expressed pharmacologically (Daou et al., 2013; Kosche et al., 2015; Long et al., 2010; Mooney & Prather, 2005). The functional forms of activation/inactivation functions and time constants were based on previous published mathematical neural models (Daou et al., 2013; Destexhe & Babloyantz, 1993; Dunmyre et al., 2011; Hodgkin & Huxley, 1952; Terman et al., 2002; Wang et al., 2003), and the parameters that were varied were merely the maximal conductances of some ionic currents that vary among the various neuronal subtypes (Daou et al., 2013), as well as the synaptic conductances. Every model neuron is represented by ordinary differential equations for the different state variables as illustrated below.

### Ion channels of model HVC neurons

We added a hyperpolarization-activated inward current conductance (I_H_) to HVC_X_ and HVC_INT_ because it is responsible for the sag seen in these neurons (Daou et al., 2013; Dutar et al., 1998; Kubota & Saito, 1991; Kubota & Taniguchi, 1998), and we added a low-threshold T-type Ca^2+^ current (I_CaT_)conductance responsible for the post-inhibitory rebound firing seen in HVC_X_ and HVC_INT_ neurons (Daou et al., 2013). A small-conductance Ca^2+^-activated K^+^ current (I_SK_) was added for HVC_RA_ and HVC_X_ neurons as it is responsible for the spike frequency adaptation feature that these two classes exhibit (Daou et al., 2013). For interneurons, we integrated a large magnitude of the delayed rectifier K^+^ current conductance allowing these neurons to undershoot the resting membrane potential as seen experimentally (Daou et al., 2013; Dutar et al., 1998; Kubota & Saito, 1991; Kubota & Taniguchi, 1998). For HVC_RA_ neurons, we added an A-type K+ current that supports the delay to spiking seen in response to depolarizing current pulses (Daou et al., 2013; Kubota & Taniguchi, 1998; Mooney & Prather, 2005). High-threshold Ca^2+^ conductance was added to all classes of HVC neurons (Daou et al., 2013; Kubota & Saito, 1991; Long et al., 2010). In total, the model was designed to include spike-producing currents (I_K_and I_Na_), a high-threshold L-type Ca^2+^ current (I_CaL_), a low-threshold T-type Ca^2+^ current (I_CaT_), a small-conductance Ca^2+^ -activated K^+^ current (I_SK_), an A-type K^+^ current (I_A_), a hyperpolarization-activated current (I_H_), and a leak current (I_L_). The membrane potential of each HVC neuron obeys the following equations:

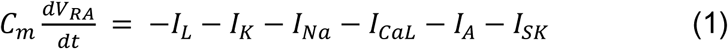

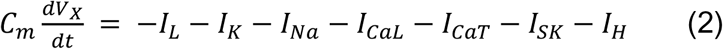

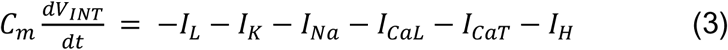

where *C*_*m*_ is the membrane capacitance. The associated equations and parameters for each of the activation/inactivation gating variables for each ionic current are given in (Daou et al., 2013) and shown below. In total, every single model HVC_RA_, HVC_X_, and HVC_INT_ neuron had a total of 6, 8 and 7 ODEs, respectively, that govern their intrinsic dynamics. Every synaptic current that was integrated to any model neuron added a new ODE to the set of ODEs governing the membrane potential of the corresponding model neuron. See APPENDIX 1 for detailed equations.

### Synaptic currents

In addition to the ionic currents above that orchestrate the internal dynamics of each HVC neuron, we integrated synaptic currents in order to reproduce the biological features of the voltage traces observed *in vivo*. Excitatory (AMPA) and inhibitory (GABA_A_) synaptic currents were used to connect neurons inside each architecture based on the pharmacological dual synaptic connections as described by Mooney and Prather (2005). Each synaptic current represents the synaptic input(s) from the presynaptic cell(s) to the particular HVC model neuron, and is modeled as *I*_𝑠𝑦𝑛_ = ∑_𝑋_ *I*_𝑋→𝑌_ where *I*_𝑋→𝑌_ = 𝑔_𝑋→𝑌_𝑠_𝑋→𝑌_(𝑉 − 𝑉_𝑋→𝑌_). Here the summation is taken over the presynaptic HVC neurons where X represents a presynaptic cell, Y represents a postsynaptic cell, 𝑉_𝑋→𝑌_ is the reversal potential for the synapse in the postsynaptic cell with 𝑉_𝑋→𝑌_ = 𝑉_𝐴𝑀𝑃𝐴_ for excitatory input and 𝑉_𝑋→𝑌_ = 𝑉_𝐺𝐴𝐵𝐴−𝐴_ for inhibitory input.

The model equations for the synaptic currents are detailed in APPENDIX 1, taken after (Destexhe et al., 1994; Varela et al., 1997). We limited our synaptic currents’ choices in all networks to AMPA (excitatory) and GABA_A_ (inhibitory) without integrating NMDA and GABA_B_ for the following reasons: 1) both AMPA and GABA_A_ currents are voltage-dependent with simple activation kinetics that do not depend on further parameters that are very hard to tune or calibrate (for example, NMDA current relies on Mg^2+^ concentration and GABA_B_ on G-proteins dynamics, both which require additional ODEs and series of parameters that we don’t know about in the HVC), 2) adding the additional synaptic currents do not have a significant contribution to the network dynamics we’re building because the emphasis is on excitation and inhibition and we could convey the mechanisms we envision that orchestrate each network with these two currents solely. Therefore, model HVC_RA_ neurons send their excitatory afferents to other HVC_RA_ neurons as well as to HVC_INT_ neurons via AMPA currents. Model HVC_INT_ neurons send their inhibitory afferents to both HVC_RA_ and HVC_X_ neurons via GABA_A_ currents. And lastly, model HVC_X_ neurons excite HVC_INT_ neurons via AMPA currents.

### Desired network activity

Our aim here is to generate the optimal and desired network activity that’s generated by the three classes of HVC neurons during singing. Therefore, we focused on model parameters and their underlying mechanisms that play key roles in 1) reproducing the patterns without breaking the sequence of activity propagation and 2) generate biologically realistic traces for each class as shown using intracellular recordings *in vivo* in a way to maintain spike shapes, burst patterns, rebound firing/bursting, subthreshold oscillations, etc … In a nutshell, the behavior of the network was considered desired and “good” if the model voltage traces for the total populations in each of the three classes of HVC neurons in the network matched the following: 1) the time-locked and characteristic bursting of HVC_RA_ neurons (3-6 spikes for a ∼10 ms duration), with spikes riding on a plateau, 2) HVC_X_ neurons eliciting 1-4 bursts (4-9 spikes per burst) that are also time-locked and that are mostly rebound bursts from inhibition (Lewicki, 1996; Mooney, 2000), 3) HVC_INT_ neurons exhibiting tonic activation with spiking and bursting throughout the song, and 4) spike frequency, spike amplitude, sags and/or rebound upon inhibition, resting membrane potential and other known features of the intrinsic properties of the three classes of HVC neurons exhibited for each of the classes that exhibit them (Daou et al., 2013).

### Maximal and synaptic conductances variations

Automated adjustment of model parameters was performed to qualitatively reproduce desired membrane potential trajectories, as described next. Fixed parameter values for HVC neurons used in the simulations are given in Table 1 in APPENDIX 2. Parameters that vary between the different model neurons are shown in Figure 2-A.

**Figure 2.**
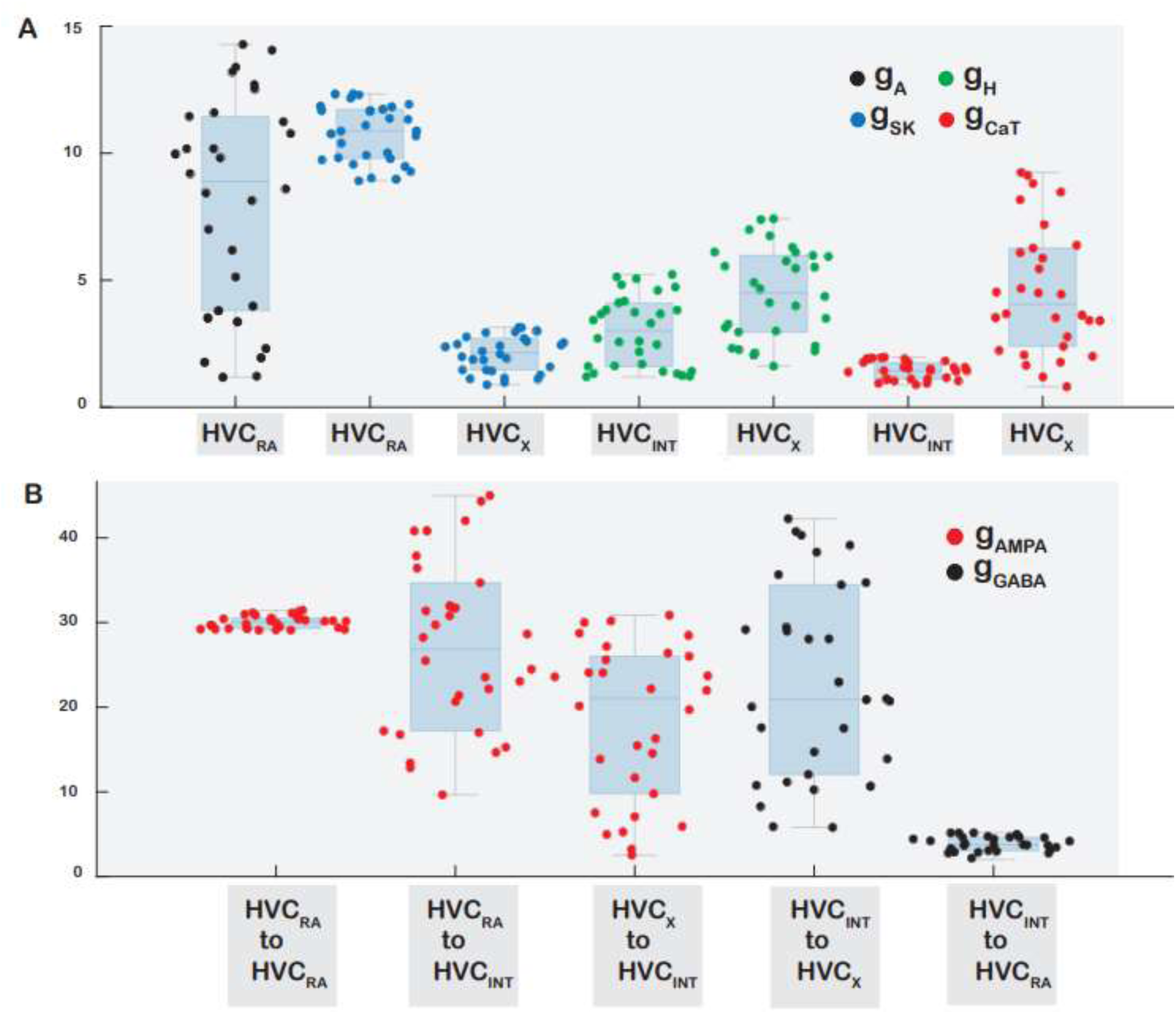
Box plots showing the ranges of ionic and synaptic currents that were allowed to vary while maintaining robust network propagation and biologically realistic in vivo behavior of all HVC neuronal classes. **A.** The ionic conductances that were varied are gA of HVC_RA_, gSK of HVC_RA_ and HVC_X_, gh and gCaT of HVC_INT_ and HVC_X_. The shown ranges reflect values whereby each neuron class is able to maintain realism in terms of electrophysiological behavior and network properties. **B.** Ranges of values of synaptic conductances that connect two classes of HVC neurons while conserving sequential activity propagation and the general network activity.

Some maximal conductances were fixed while others were allowed to vary. We fixed g_Na_and g_K_ for each class of the HVC neurons to values that had been shown earlier to accurately fit the spike morphologies (upstrokes and downstrokes of action potentials, plateaus, etc…) in response to applied current given *in vitro* (Daou et al., 2013). For example, HVC_INT_ neurons’ spikes exhibit a relatively large undershoot of the resting membrane potential, while HVC_X_ and HVC_RA_ spikes ride on a plateau with characteristic properties (Daou et al., 2013). We also fixed the value of g_CaL_because we could achieve the same accuracy of fitting by varying g_SK_, and so could not distinguish between the two.

The four key conductances in our model that played crucial roles in controlling not only the intrinsic properties of the HVC neurons they’re expressed in but also in shaping overall network activity and sequence propagation are g_SK_, g_h_, g_A_ and g_CaT_. As a recap, g_SK_ is expressed in HVC_X_ and HVC_RA_, g_H_ and g_CaT_ in HVC_X_ and HVC_INT_, and g_A_ in HVC_RA_ only (Daou et al., 2013). Random variations in these four parameters were performed to qualitatively reproduce membrane potential trajectories of the three classes of HVC neurons as seen firing when the bird is singing.

In a previous study, Daou and Margoliash (2020) showed that intracellular recordings from X-projecting neurons in adult zebra finch brain slices share similar spike waveform morphologies, with modeling indicating similar magnitudes of their principal ion currents. To that end, we fixed in our network the intrinsic properties of the population of HVC_X_ neurons to the same values. Therefore, the automated variations in the conductances held for HVC_X_ neurons were done at the population level by varying a corresponding ionic conductance value (say, g_SK_) and setting its value to all of the HVC_X_ population. We also checked the effects of removing this constraint in Figure 13, that is, allowing the intrinsic properties of HVC_X_ neurons to vary like other parameters.

We first manually selected the four key conductances to default values that generate the desired behavior of the network as described in the previous section. Network robustness to varied maximal conductances as well as the legitimate ranges for each maximal conductance was determined by simulating the network many times, each time with a random variation in the maximal conductance about its default value. The network response was considered accurate if the network generated sequential bursting and all of the desired features described earlier. The range of variation of the randomly-varying parameter was increased or decreased randomly up until the point that the network ceased to be accurate.

The variation in parameters was done at the population level (setting the maximal conductance for all neurons of the same class to a single value and randomly changing that value for all), well as on the individual neuronal level (varying the maximal conductance for one neuron at a time of the same class while fixing the others to their default values). This is different for the HVC_X_ parameters, where the corresponding conductances at the population and the neuronal levels are considered the same since we assumed same intrinsic properties as described earlier. If any of the simulations generated networks where any of the HVC_RA_ neurons elicit bursts exhibiting spikes outside the 3-6 number of spikes range or duration of any burst longer than 10 ms, then we ignore that parameter value. Similarly, if any of the simulations generate networks where any of the HVC_X_ neurons eliciting more than 4 bursts, individual bursts exhibiting spikes outside the 4-9 spikes/burst range, then the corresponding parameters are ignored. Moreover, we also ignore parameters that generate unrealistic intrinsic properties of individual HVC neurons; for instance, if the resting membrane potentials of individual HVC neurons were outside the reported ranges (Daou et al., 2013), if HVC_X_ and HVC_INT_ neurons failed to generate sags and rebounds (depolarization or bursting) in response to inhibition, or if spikes’ amplitudes were not realistic (HVC_X_ and HVC_RA_ spikes not riding on plateaus or HVC_INT_ spikes not undershooting the RMP). We realize this might be inducing tough constraints on the selection of the parameters limiting the space for which the conductances are allowed to vary, but we opted for this method in order to generate the optimal ranges in which intrinsic and synaptic conductances are able to reproduce the biophysically realistic firing patterns seen during singing. We therefore ended up with lists of ranges for each conductance that was varied and in each class of HVC neuron, such that the desired network activity is maintained. Figure 2-A shows the ranges for each maximal conductance that had been allowed to vary for the three classes of HVC neurons, while maintaining robust network propagation and biologically realistic in vivo behavior.

Similar to what was done with the maximal ionic conductances, we conducted automated and random variations for all synaptic conductances in the model (no synaptic conductance was fixed). Figure 2-B reports the ranges of the synaptic conductances that were able to maintain the robustness of network propagation and the general *in vivo*-like desired behavior of all neuronal classes. All synaptic conductances showed considerable ranges during which sequential activity is propagated and the overall desired network activity is maintained, with the exception of the GABA conductance from HVC_INT_ to HVC_RA_ (𝑔_𝐺𝐴𝐵𝐴*I*𝑁𝑇→𝑅𝐴_). Increasing 𝑔_𝐺𝐴𝐵𝐴*I*𝑁𝑇→𝑅𝐴_ to larger magnitudes would induce an inhibition in the HVC_RA_ pushing its voltage below its resting membrane potential (due to the dense bursting and firing in HVC_INT_), and this is not realistic because it’s been shown that during singing HVC_RA_ neurons ride on a depolarizing plateau throughout the song (Long et al., 2010). Moreover, while the intrinsic properties of HVC_X_ neurons were set to the same values, the synaptic parameters associated with each HVC_X_ neuron (afferent and efferent) were allowed to vary from one neuron to another.

Moreover, to account for synaptic variability, we introduced a stochastic input current of the form *I*_𝑛𝑜𝑖𝑠𝑒_(𝑡) = 𝜎. 𝜉(𝑡) where 𝜉(𝑡) is a Gaussian white noise with zero mean and unit variance, and 𝜎 is the noise amplitude. This stochastic drive was introduced to every model neuron and it mimics the fluctuations in synaptic input arising from random presynaptic activity and background noise. For values of 𝜎 within 1-5% of the mean synaptic conductance, the stochastic current has no effect on network propagation. For larger values of 𝜎, the desired network activity was disrupted or halted.

## Results

Adult zebra finches generate intricate songs composed of sequences of distinct song elements, each characterized by a stereotypical acoustic pattern across every rendition of song. The neural circuitry that governs this behavior consists of HVC_RA_ neuronal population where each emits a single and stereotyped 6-10 ms burst during each rendition of song, HVC_X_ neurons eliciting 1-4 bursts that are similarly time locked to vocalizations, and HVC_INT_ neurons that tend to burst densely throughout song. Intrinsic and synaptic mechanisms that orchestrate these neurons’ behaviors are well known (Daou et al., 2013; Kornfeld et al., 2017; Mooney & Prather, 2005). We next describe the steps of building our biophysical network model to describe this ongoing behavior in the following order: 1) tuning the synaptic parameters to fit the dual-intracellular recording traces collected experimentally by Mooney and Prather (2005) as well as Kosche et al (2015), and then 2) describing the network components that are essential for the patterned output of the system, as well as the internal dynamics of the network that govern the strength and duration of individual bursts, the duration of silent gaps between bursts, sparseness versus tonicity, the interplay between excitation and inhibition, role of intrinsic properties and so on that explain how the firing activity of the three classes of HVC neurons propagate through the network in sequential manner.

### Tuning synaptic parameters

We initiated our HVC network modeling study calibrating the synaptic parameters (excitatory and inhibitory currents’ activation/inactivation constants, etc..) by reproducing the voltage traces elicited by the dual intracellular recordings from identified pairs of HVC neurons in brain slices conducted by Mooney and Prather (2005). While we are using off-the-shelf synaptic currents from the literature (Destexhe et al., 1994; Varela et al., 1997), we needed to make sure that the synaptic parameters used can replicate the dual synaptic connectivity patterns (strengths of excitation/inhibition, magnitudes of voltage deflections and other trace morphologies). Mooney and Prather’s findings revealed robust disynaptic feedforward inhibition from HVC_RA_ to HVC_X_ neurons (mediated by HVC_INT_ neurons), potent monosynaptic excitation from HVC_RA_ and HVC_X_ to HVC_INT_ neurons (via NMDA and AMPA currents), and substantial monosynaptic inhibition from HVC_INT_ neurons to HVC_RA_ and HVC_X_ (via GABA currents).

Figure 3 displays the dual intracellular recordings conducted by Mooney and Prather (left column) as well as the mathematical model replications (right column) after the synaptic parameters’ calibration. DC-evoked action potentials in HVC_RA_ neurons trigger inhibitory postsynaptic potentials (iPSPs) in HVC_X_ neurons (Fig. 3-A1), as well as fast depolarizing postsynaptic potentials (dPSP) in HVC_INT_ neurons (Fig. 3-B1). To replicate the effects of stimulating HVC_RA_ neurons onto the other two classes of HVC neurons, we connected one HVC_RA_ neuron to excite one HVC_INT_ neuron via an AMPA current. The HVC_INT_ neuron in its turn was connected with one HVC_X_ neuron via a GABA current, thereby making a di-synaptic pathway from HVC_RA_ to HVC_X_. DC-evoked action potentials in the model HVC_RA_ neuron (brief ∼10 ms depolarizing current pulses (0.5 nA, similar to what Mooney and Prather applied to HVC_RA_ neurons) evoked a fast-depolarizing postsynaptic potential (dPSP) in the corresponding model HVC_INT_ neuron (Fig. 3-B2) as well as inhibitory postsynaptic potential (iPSP) in the corresponding model HVC_X_ neuron (Fig. 3-A2) mediated via HVC_INT_. Similarly, DC-evoked action potentials in HVC_INT_ neurons generate fast iPSPs in HVC_X_ neurons (Figures 3-C1, D1). The unequal magnitude of the four sags elicited in the HVC_X_ neuron (due to the four brief stimuli) and the large sag after the stimuli ends (Fig. 3-C1), as well as the jagged long sag in response to the repetitive action potentials elicited by the HVC_INT_ neuron (Fig. 3-D1) is probably due to the fact that the corresponding HVC_X_ neurons are receiving multiple synaptic inputs from neurons other than the neurons being stimulated by Mooney and Prather, which adds to the underlying nonlinearities of the responses being recorded. Model HVC_INT_ and HVC_X_ neurons were connected and synaptic parameters were calibrated to generate similar waveforms by giving brief ∼10 ms depolarizing current pulses of 0.5 nA to HVC_INT_ (Fig. 3-C2), or giving a DC-pulse of 0.5 nA for 500 ms (Fig. 3-D2). In the model, the sag seen in the HVC_X_ neuron response is due to the build-up of the H-current as the model HVC_INT_ neurons continues firing, exerting its inhibition onto the HVC_X_ neuron. Finally, model HVC_X_-HVC_INT_ monosynaptic connectivity (Fig. 3-E2) was calibrated to match the experimental findings (Fig. 3-E1).

**Figure 3.**
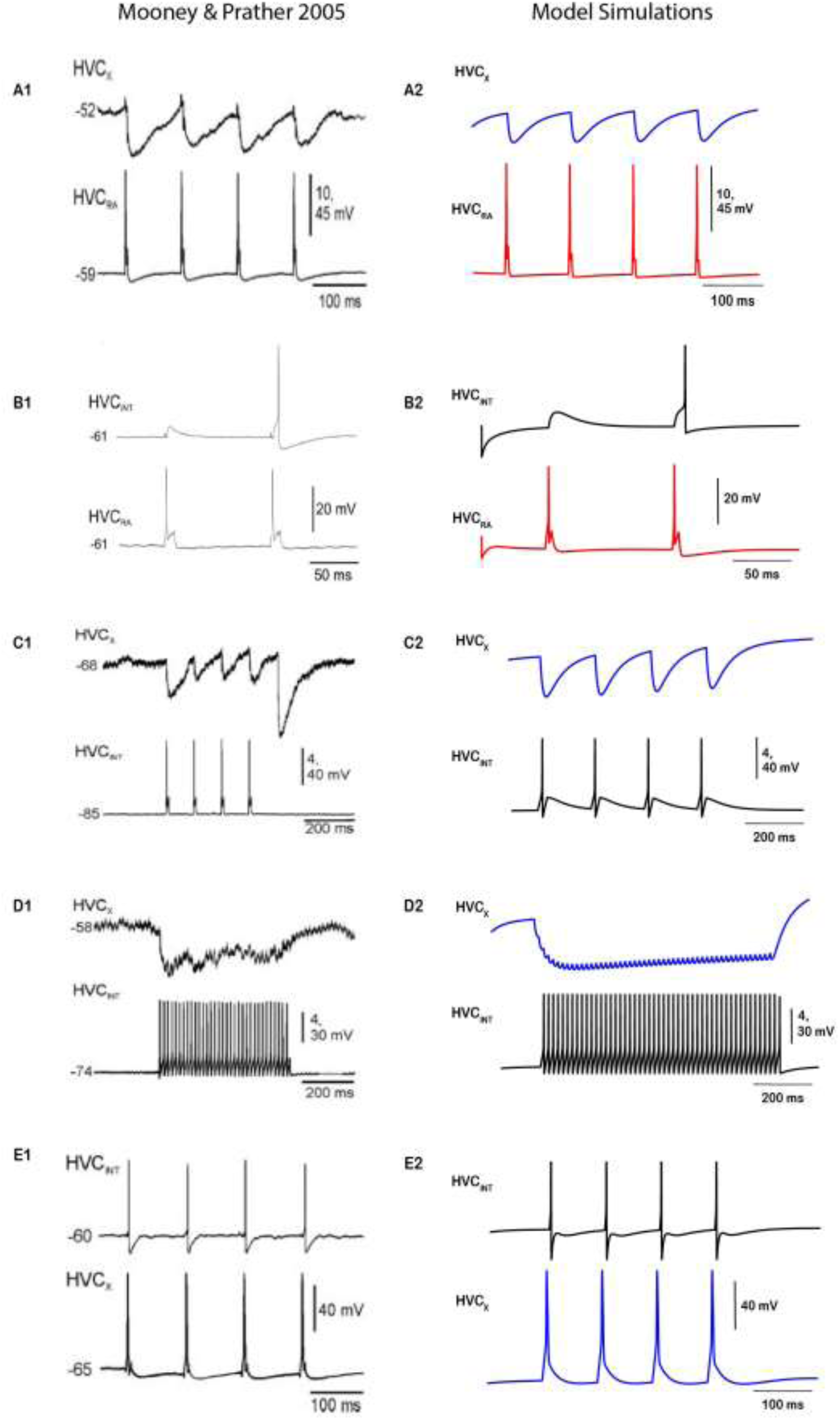
Model output compared to experimental results obtained by Mooney and Prather (2005). DC-evoked action potentials in HVC_RA_ neurons trigger iPSPs in HVC_X_ neurons (**A1**) as well as fast dPSPs in HVC_INT_ neurons (**B1**). Brief (∼10 ms) depolarizing current pulses (0.5 nA) applied to model HVC_RA_ neuron (same values used as by Mooney and Prather (2005)) evokes similar responses in the corresponding model HVC_X_ (**A2**) and model HVC_INT_ (**B2**) neurons. DC-evoked action potentials in HVC_INT_ neurons generate fast iPSPs in HVC_X_ neurons (**C1**, **D1**). Similar responses were elicited in model HVC_X_ neurons when model HVC_INT_ neuron was stimulated by brief (10 ms) depolarizing pulses (0.5nA) (**C2**) or when it was given a DC-pulse of 0.5 nA for 500 ms (**D2**). Finally, HVC_INT_ neurons elicit fast dPSPs when HVC_X_ neurons are injected with 10 ms pulses of 0.5 nA current (**E1**), which was simulated in the model (**E2**). In this and subsequent figures (unless otherwise specified), HVC_X_ neurons’ traces are represented in blue, HVC_RA_ neurons in red, and HVC_INT_ neurons in black. Panels in the left column adopted from Mooney and Prather (2005).

### Network Architecture: Non-sequential random sampling in HVC

We next describe the activity patterns generated by our network model, and explain how the firing activity of the three classes of HVC neurons propagate through the network in sequential bursts of activity. The network developed is composed of chains of HVC subnetworks or microcircuits, each with its own intrinsic dynamics. The microcircuit represents a basic architectural unit that encodes for a syllable or a sub-syllabic segment (SSS) in the motif (Fig. 4). Each neuron in a microcircuit is representative of a neural population. In other words, a model neuron (belonging to any class) firing is representative of a population of that neuronal class firing, which could be many neurons of the same class exhibiting very similar intrinsic and synaptic properties leading to their firing at the same time. We refer here to “microcircuits” in a more functional sense, rather than rigid, isolated spatial divisions (Cannon et al. 2015). A microcircuit in our model reflects the local rules that govern the interaction between all HVC neuron classes within the broader network, and that are essential for proper activity propagation. The number of microcircuits in the chain determines the number of SSSs in the motif. We envision the HVC to be composed of many copies of such microcircuit chains that are associated with SSSs with roughly synchronized activity. The duration of the sub-syllabic segments need not be the same; therefore, the number of neurons that each microcircuit encompasses need not be equal as we will see next. Moreover, while silent gaps are integral to the overall process of song production, we have not elaborated on them in this model due to the lack of a clear, biophysically grounded representation for the gaps themselves at the level of HVC. Our primary focus here is on modeling the active, syllable-producing phases of the song, where the HVC network’s sequential dynamics are critical for song. However, one can think the encoding of silent gaps via similar mechanisms that encode SSSs, where each gap is encoded by similar microcircuits comprised of the three classes of HVC neurons (let’s call them GAP, rather than SSS) that are active only during the silent gaps. In this case, the propagation of sequential activity is carried throughout the GAPs from the last SSS of the previous syllable to the first SSS of the subsequent syllable.

**Figure 4.**
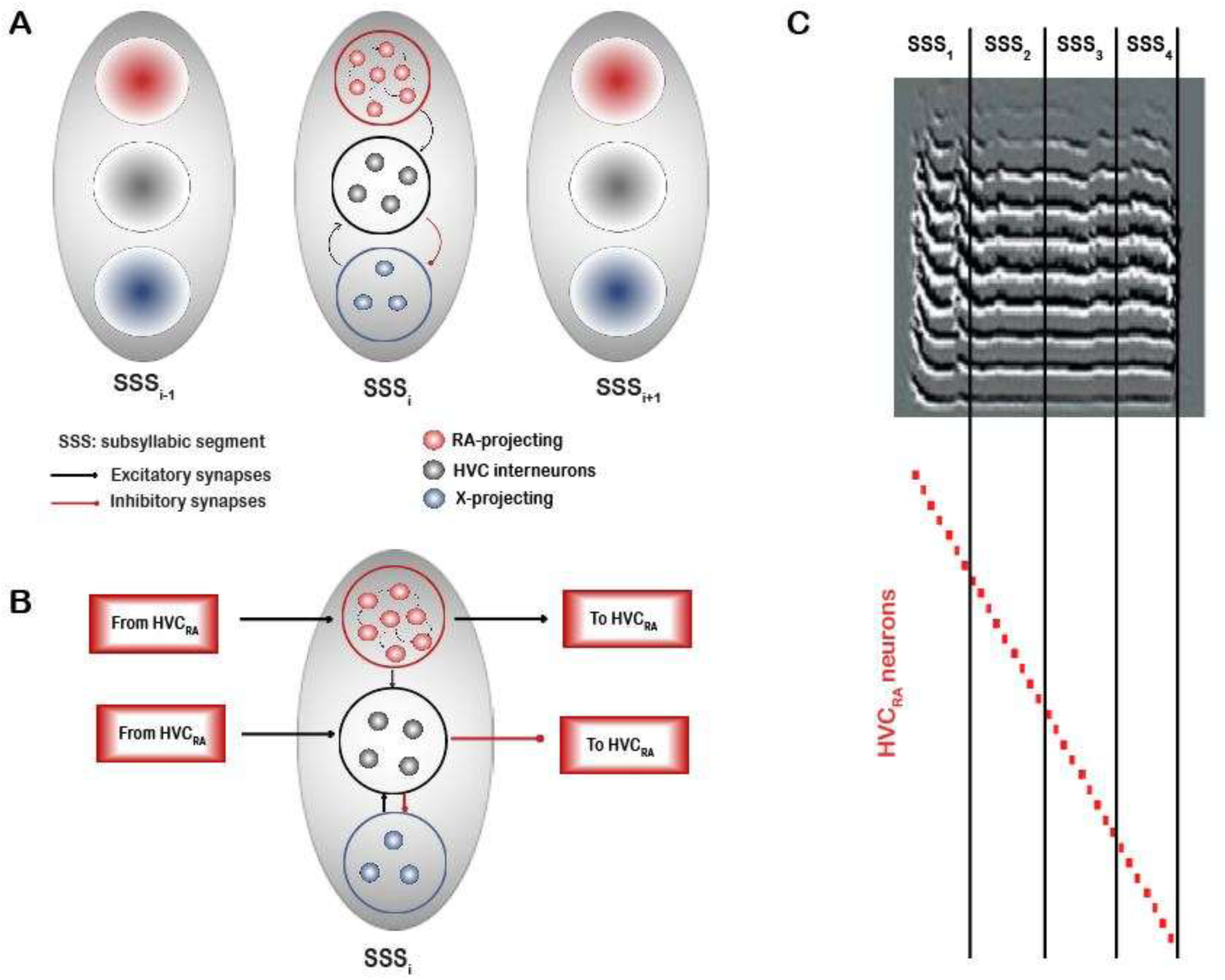
Cartoon diagram illustrating the network architecture configuration. **A.** Each grey oval represents a microcircuit encoding for a sub-syllabic segment (SSS). The number of microcircuits is envisioned to be equal to the number of SSSs representing the song. Each microcircuit contains a number of HVC_RA_, HVC_X_ and HVC_INT_ neurons selected randomly from the total pool of neurons (see text). **B**. Within each microcircuit, HVC_RA_ neurons are connected to each other in a chain-like mode and they send excitatory afferents to HVC_INT_ neurons in the same and other microcircuits, selected randomly. HVC_INT_ neurons send GABAergic synapses onto HVC_X_ neurons in the same microcircuit only as well as to HVC_RA_ neurons in any other random microcircuit except the microcircuit they belong to. Finally, HVC_X_ neurons send excitatory afferents to HVC_INT_ neurons in the same microcircuit. Activity starts by a small DC pulse to the first HVC_RA_ neuron in the first microcircuit. Activity propagates from one microcircuit to another by excitatory coupling between the last HVC_RA_ neuron in microcircuit i and the first HVC_RA_ neuron in microcircuit i+1. **C.** During singing, the propagation of activity unfolds across the chain of microcircuits, such that neurons belonging to microcircuit x gets activated and encode for SSS_x_.

The network is comprised of a total pool of HVC_RA_ neurons (120 neurons, red circles), a pool of interneurons (50 neurons, black circles) and a pool of HVC_X_ neurons (50 neurons, blue circles), thereby maintaining a 2:1:1 proportionality factor across the populations of HVC_RA_: HVC_INT_: HVC_X_ as reported earlier (Kornfeld et al., 2017; Wild et al., 2005). The total number of neurons in the pool is arbitrary and can be made larger, but we limited it to these values given the huge number of ODEs that are being simulated (∼2000 ODEs). Figure 4-A shows the network diagram illustrating three sample microcircuits encoding for three sub-syllabic segments (SSS_i-1_, SSS_i_, and SSS_i+1_) in sequence. Each microcircuit (enclosed by a gray oval) is made up of a number of HVC_RA_ neurons (red circles) and a number of HVC_INT_ neurons (black circles), all selected randomly from their corresponding pools, as we will describe next. In our model, we limited the number of microcircuits to twenty (i.e., the motif is encoded by twenty sub-syllabic segments), but this number is also arbitrary and can be made larger or smaller.

### Network organization

The total pool of HVC_RA_ neurons is comprised of smaller groups of HVC_RA_ neurons, the number in each group of which is chosen randomly from the pool (red background circles in each gray oval). Each group of HVC_RA_ neurons belongs to a unique microcircuit and no HVC_RA_ neuron is allowed to be part of more than one microcircuit for reasons described next. Since we set the motif to be represented by twenty microcircuits in our illustration, HVC_RA_ neurons were recruited to their corresponding microcircuits randomly, with each microcircuit allowing a random number (3 – 10) neurons of the RA-projectors to belong to it. In addition to that, the numbers of HVC_INT_ neurons (black background circles) that a microcircuit exhibit is a random number between 1 and 4, as well as the number of HVC_X_ neurons (blue background circles) in each microcircuit is random between 1 and 4, where the individual neurons are selected arbitrarily one neuron at a time from their corresponding pools. Each HVC_INT_ neuron can belong to a single microcircuit and similarly each HVC_X_ neuron can belong to a single microcircuit for reasons described next.

### Synaptic Connectivity

Within each microcircuit, HVC_RA_ neurons are selected randomly, one after the other, to send AMPA excitatory synapses to each other in a chain-like mode. Specifically, if there are *m* HVC_RA_ neurons recruited to belong to microcircuit 𝑖 (where *m* is the random number generated between 3 and 10 in this case), a neuron from the *m* is first selected randomly and designated as the first neuron in the chain (*HVC^i^_RA_1__*). After that, a second neuron 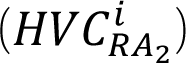 from the remaining *m* − 1 is selected randomly and an AMPA synapse is connected from 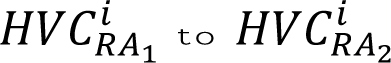. Similarly, a third neuron 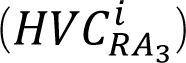 is selected randomly from the remaining *m* − 2 neurons and an AMPA synapse is connected from 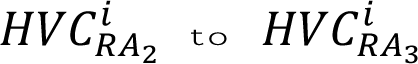 and so on, until the HVC_RA_ neurons in every microcircuit are connected together (Fig. 4A, small red circles in each microcircuit).

Each HVC_INT_ in a microcircuit was assigned a random number (between 3 and 8) of excitatory AMPA connections from the HVC_RA_ neurons in the same microcircuit it belongs to, as well as from HVC_RA_ neurons in the other microcircuits (Fig. 4B). In their turn, each HVC_INT_ neuron sends a random number (between 2 and 4) of GABAergic inputs to HVC_RA_ neurons, chosen arbitrarily from any microcircuit in the chain except the microcircuit that the HVC_INT_ neuron belongs to, due to the following reason: if HVC_INT_ inhibits HVC_RA_ in the same microcircuit, some of the HVC_RA_ bursts in the microcircuit might be silenced by the dense and strong HVC_INT_ inhibition breaking the chain of activity. However, if HVC_INT_ inhibits HVC_RA_ in any other microcircuit, activity is ensured to propagate because the HVC_INT_ inhibition of the corresponding HVC_RA_ would arrive at times that are not the “assigned” times of the HVC_RA_ to elicit their ultra-sparse code (Hahnloser et al., 2002).

Unlike HVC_RA_, HVC_INT_ neurons belonging to a particular microcircuit can burst at times other than the moments when the corresponding encoded SSS is being “sung”; however, we chose to house interneurons within microcircuits for the mere fact that any given interneuron cannot inhibit any given HVC_RA_ neuron, rather there are some “rules of engagement” where we ensure that no inhibition arrives to any HVC_RA_ neuron while it’s eliciting its burst of activity (Fig. 4B). In other words, what makes a particular interneuron belong to this microcircuit or the other is merely the fact that it cannot inhibit HVC_RA_ neurons that are housed in the microcircuit it belongs to for the reasons described. In this regard, this arrangement is similar to the Cannon et al (2015) model in the context of structured inhibition amid the ongoing feedforward excitation.

At the HVC_X_ side, each X-projecting neuron excites via AMPA currents a random selection (1 - 3) of HVC_INT_ neurons that belong to the same microcircuit, and in their turn, each HVC_INT_ neuron inhibits via GABA synapses a random selection (1 - 2) of HVC_X_ neurons in the same microcircuit (Fig. 4B). These numbers are again arbitrary, but we limited the number of connections from HVC_INT_ to HVC_X_ due to that fact that X-projecting neurons in our model fire upon rebound from inhibition, and the more inhibitory inputs they receive, the more rebound bursts they elicit, which is not realistic since HVC_X_ neurons are known to elicit 1 - 4 bursts during singing (Fujimoto et al., 2011; Kozhevnikov & Fee, 2007), which is what we achieved with this number of synapses. HVC_X_ neurons were selected to be housed within microcircuits and their synapses connecting to interneurons within the same microcircuit due to the following reason: if an HVC_X_ neuron belonging to microcircuit 𝑖 sends excitatory input to an HVC_INT_ neuron in microcircuit 𝑗, and that interneuron happens to select an HVC_RA_ neuron from microcircuit 𝑖 as its afferent inhibitory connection (via random sampling), then the propagation of sequential activity will halt, and we’ll be in a scenario similar to what was described earlier for HVC_INT_ neurons inhibiting HVC_RA_ neurons in the same microcircuit. Similarly, if an HVC_INT_ neuron in microcircuit 𝑖 inhibits an HVC_X_ neuron in another microcircuit 𝑗, and that HVC_X_ neuron excites an interneuron that synapses onto an HVC_RA_ from microcircuit 𝑖, then sequential activity might be disrupted.

While HVC_RA_ neurons are connected to each other in each microcircuit in a chain-like mode as described earlier, the microcircuits interact with each other via the projections from the last HVC_RA_ in a microcircuit 𝑖 to the first HVC_RA_ in a following microcircuit 𝑖 + 1. The network is kick-started by a stochastic DC input to 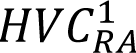, that is, only 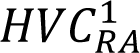 receives input from outside HVC. The propagation of sequential activity along with the realistic firing of the three classes of HVC neurons is maintained and orchestrated by HVC’s intrinsic and synaptic processes without relying on extrinsic inputs as shown next (Figures. 5-11).

**Figure 5.**
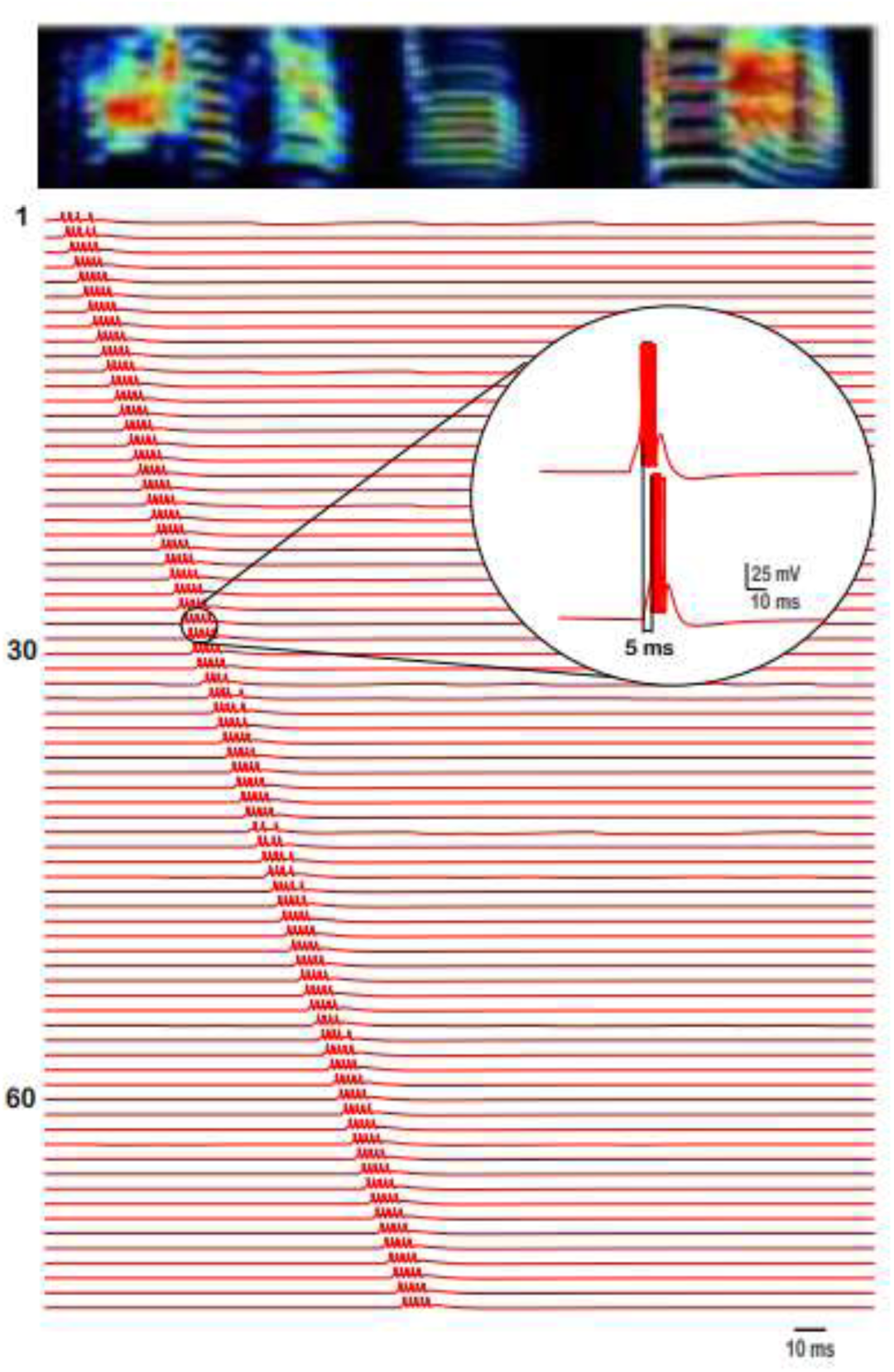
Spiking patterns of 120 HVC_RA_ neurons (labeled with numbers) showing the propagation of sequential activity. The neural traces are aligned by the acoustic elements of a spectrogram from an exemplar bird’s song illustrating the firing of HVC_RA_ neurons with respect to ongoing part of a song. The inset shows a zoomed version of two subsequent HVC_RA_ neurons firing patterns illustrating the delay between their individual bursts.

### Sequential propagation of HVC_RA_ activity

The activity patterns that RA-projecting neurons of this network display are illustrated in Figure 5. HVC_RA_ neurons burst extremely sparsely generating at most a single burst per simulation (song motif) and with different HVC_RA_ neurons bursting at different times in the song. HVC_RA_ bursts had a duration of 8.12 ± 0.89 ms, and comprised of 4.76 ± 0.48 spikes. The inset of Fig. 5 shows the delay from the onset of the first spike in an HVC_RA_ neuron to the onset of the first spike in the next HVC_RA_ it synapses to. This duration is controlled by two factors: 1) the magnitude of the AMPA conductance, with larger magnitudes corresponding to shorter delays and vice versa (Fig. 6A). In short, the faster the AMPA current is (modeled as larger magnitude in the 𝑔_𝐴𝑀𝑃𝐴_ parameter that connects two HVC_RA_ neurons), the shorter the delay between the successive HVC_RA_ bursts. 2) the magnitude of the A-type K^+^ current conductance (𝑔_𝐴_) with larger magnitudes corresponding to longer delays (Fig. 6B). This conductance increases rapidly on depolarization due to *I*_𝐴_’s fast activation. The rapid increase in 𝑔_𝐴_ halts after a few milliseconds and switches to a slow decrease that is due to the slow inactivation that this current exhibit. The slow decrease is reflected in the voltage trace as a slow depolarization in the membrane potential (encoding for the delay to spiking), and this allows the model HVC_RA_ neuron to eventually escape the inhibition produced by *I*_𝐴_ and fire its delayed burst.

**Figure 6.**
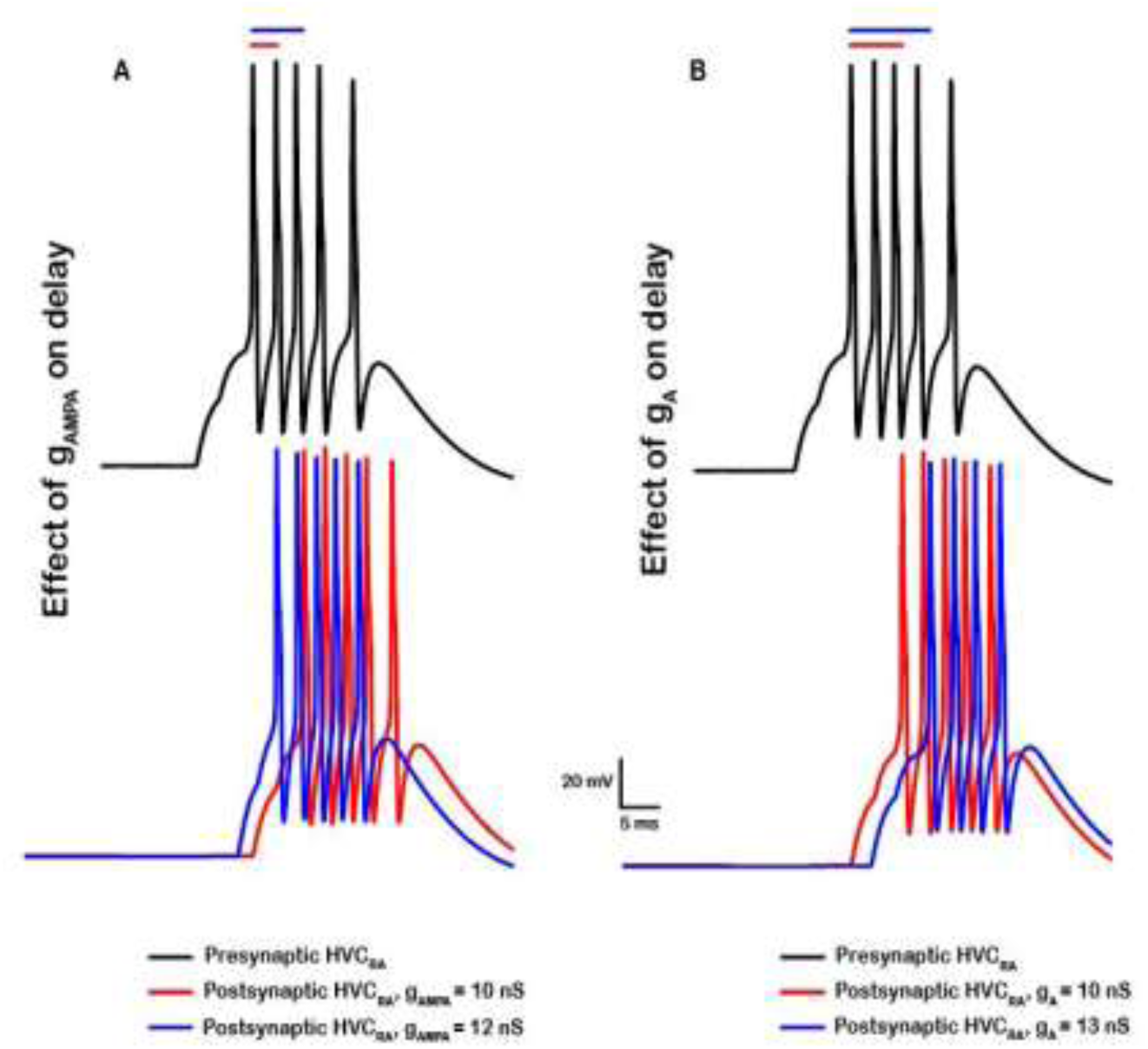
Effects of the AMPA synaptic conductance and A-type K^+^ conductance on the delay between two successive HVC_RA_ bursts. **A**. Presynaptic model HVC_RA_ neuron (top, black) is connected to a postsynaptic model HVC_RA_ neuron and the corresponding AMPA excitatory conductance (𝑔_𝐴𝑀𝑃𝐴_) was increased from 10 nS (bottom, red) to 12 nS (bottom, blue), while keeping all other parameters fixed. Increasing 𝑔_𝐴𝑀𝑃𝐴_ reduce the delay between the peaks of the pre- and post-HVC_RA_’s first spikes and increase the number of spikes in the postsynaptic neuron. **B**. Larger magnitudes of the A-type K^+^ conductance (𝑔_𝐴_) leads to longer delays to spiking. While keeping all intrinsic and synaptic parameters fixed (𝑔_𝐴𝑀𝑃𝐴_ = 10), increasing 𝑔_𝐴_ from 10 nS (bottom, red) to 13 nS (bottom, blue) delayed the onset to spiking and reduced the number of spikes. Bars on the top shows the duration in ms between the peak of first action potential in the presynaptic neuron to the peak of the first action potential in the postsynaptic neuron.

The number of spikes in HVC_RA_ neurons and the strength of the burst is controlled by three factors: 1) the strength of the AMPA conductance itself where stronger excitatory coupling corresponds to larger number of spikes (Fig. 6A), 2) the A-type K^+^ conductance where larger magnitudes of the conductance reduce the general excitability of the HVC_RA_ neuron and limit its number of spikes (Fig. 6B), and 3) the interplay between the L-type Ca^2+^ (𝑔_*C*𝑎𝐿_) and the Ca^2+^-dependent K^+^ (𝑔_𝑆𝐾_) conductances that control the strong adaptation which these neurons exhibit (Daou et al., 2013). In short, intracellular Ca^2+^ (due to *I*_*C*𝑎𝐿_) accumulates during the HVC_RA_ burst. This result in a buildup of Ca^2+^-activated K^+^ current (*I*_𝑆𝐾_) that terminates the HVC_RA_ burst, which in turn terminates any burst in a post-synaptic neuron since HVC_RA_ can no longer provide excitation.

The propagation of sequential bursting in HVC_RA_ neurons halts and the chain of HVC_RA_ activity is broken if any of the following is satisfied: 1) if an AMPA conductance for any of the HVC_RA_’s that connects it to the next HVC_RA_ in its chain is small enough such that it’s not able to elicit sufficient excitability on the postsynaptic side, 2) if a 𝑔_𝐴_ conductance (Fig. 7A) or a 𝑔_𝑆𝐾_ conductance (Fig. 7B) in any of the HVC_RA_’s is large enough (mimicking an up-regulation in the corresponding channel) to eliminate the corresponding HVC_RA_’s burst. Therefore, in order to generate accurate HVC_RA_ bursting patterns that maintain the sequential propagation of neural activity, accurate number of spikes and delays between spikes across the population of HVC_RA_’s in all microcircuits, as well as the general intrinsic properties of the HVC_RA_ neurons’ spike morphologies (see Methods), both intrinsic and synaptic constraints are needed with key roles in this process for the A-type K^+^ and the Ca^2+^-dependent K^+^ currents that HVC_RA_ model neurons exhibit, as well as the AMPA currents connecting the HVC_RA_ population together within and across microcircuits. Recall that we envision each HVC neuron of any class in our model as a representative of a neural population of the same class that exhibits the same intrinsic as well as afferent and efferent synaptic connectivity. Therefore, in Fig. 7 and the subsequent figures (10,12 and 13) where we show disruption of sequential activity due to changes in synaptic or intrinsic properties of HVC neurons, the modeled synaptic/intrinsic changes to the selected neurons shown are envisioned to be changes applied to the whole neural population encoded by our model neuron. In other words, disrupting the properties of a single neuron within that neural population will not cause harm to the propagation of activity due to what could be thought of as homeostatic mechanisms of the network (Golowasch et al., 1999; Marder & Goaillard, 2006; Williams et al., 2013). This redundancy within the population allows the propagation of activity to be maintained. It is an important feature of our model and is consistent with biological observations where neural populations exhibit robust collective behavior and the loss of a single neuron does not result in a major disruption of network activity.

**Figure 7.**
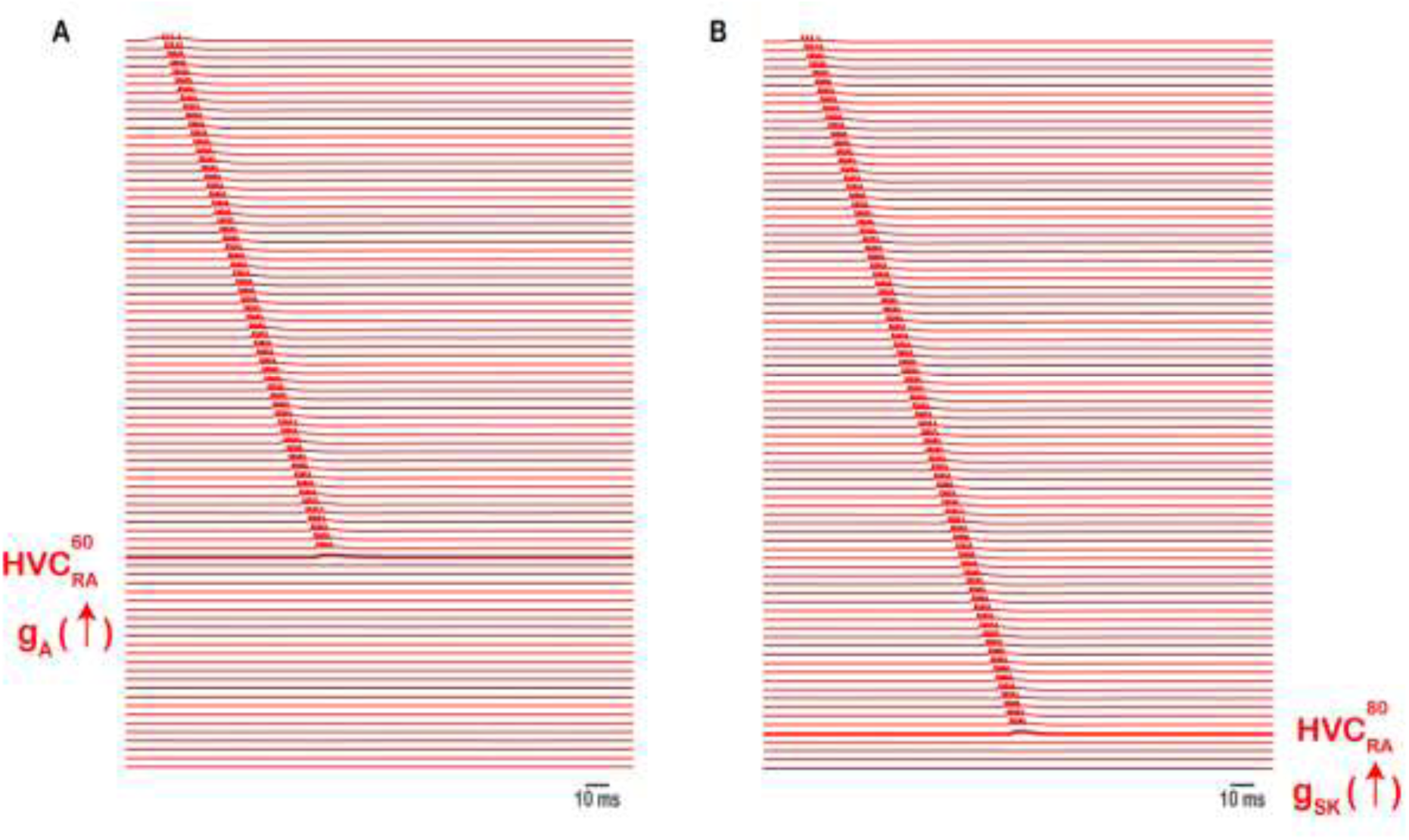
Intrinsic changes in HVC_RA_ halts the propagation of sequential activity. Up-regulating the A-type K^+^ current (**A**) or the Ca^2+^ - dependent K^+^ current (**B**) in exemplar neurons 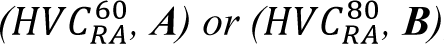, by increasing 𝑔_𝐴_ 10-fold (**A**), or 𝑔_𝑆𝐾_from 15-fold nS (**B**), reduces the excitability of corresponding HVC_RA_ neuron markedly, eliminating its corresponding burst and breaking the sequence.

### Bursting patterns of HVC_INT_ and HVC_X_ neurons

The activity patterns that HVC_INT_ neurons exhibit in our network architecture are illustrated in Figure 8A (shown here for 10 HVC_INT_ neurons). In contrast to the sparse activity in RA-projecting neurons, HVC interneurons generate multiple bursts and spike densely as reported during the song (Hahnloser et al., 2002). The number of bursts each HVC_INT_ neuron exhibits as well as the strength of each burst is controlled by network and synaptic mechanisms as described next (Figures. 9, 10). HVC_X_ neurons in their turn generate 1 - 4 bursts per song motif (Fig. 8B, shown here for 10 HVC_X_ neurons) similar to experimental results (Fujimoto et al., 2011; Hahnloser et al., 2002). Similarly, the number of bursts and strength of each HVC_X_ burst depends on a set of synaptic and intrinsic mechanisms, illustrated in Figures 11 and 12. HVC_X_ neurons differed in the number of bursts and the number of spikes per burst rendering the results more realistic.

**Figure 8.**
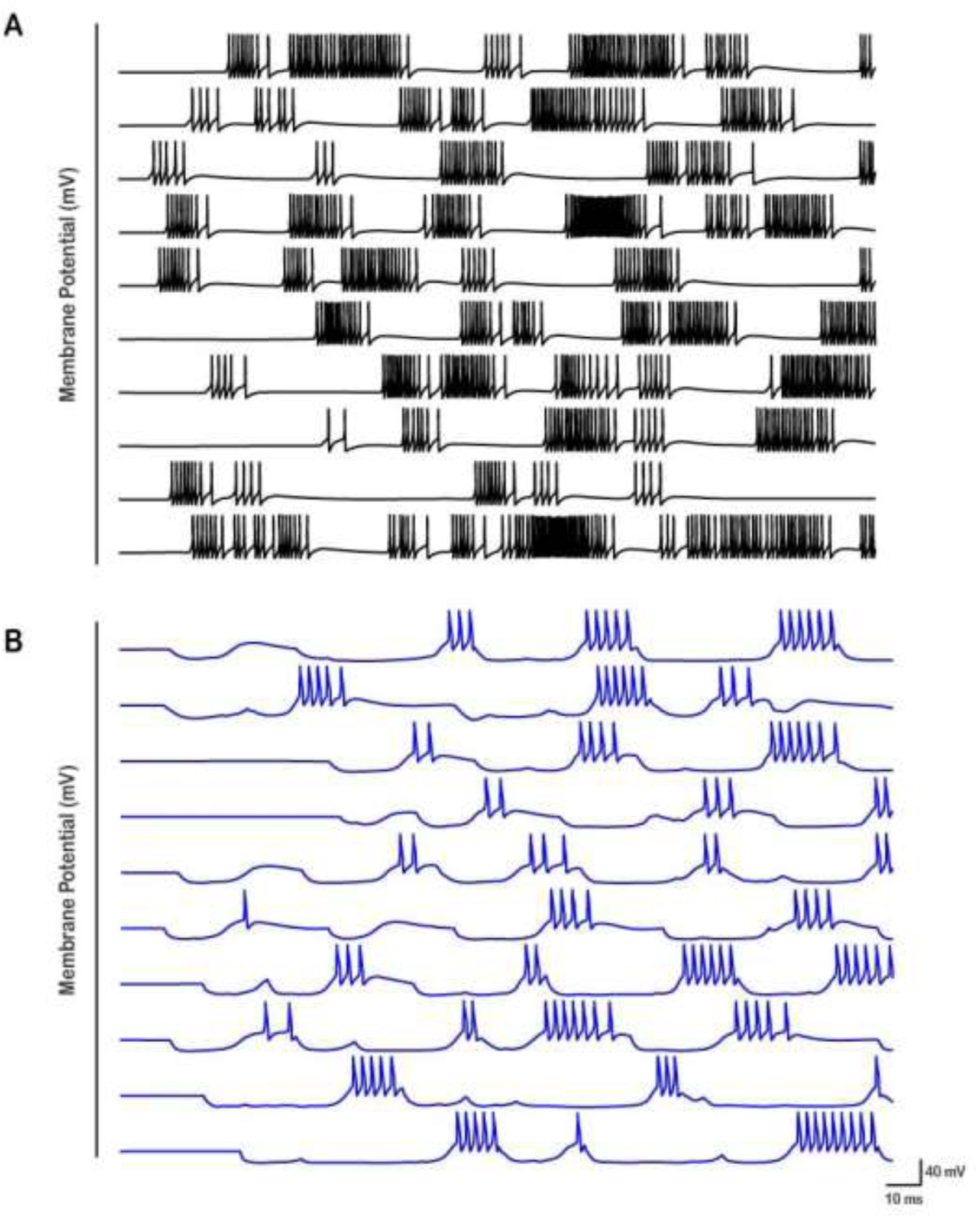
Activity patterns for 10 HVC_INT_ and 10 HVC_X_ neurons are illustrated. **A**. HVC interneurons displays dense spiking and bursting throughout the song, due to the dense HVC_RA_ – HVC_INT_ and HVC_X_ – HVC_INT_ excitatory coupling (Fig. 4). **B**. HVC_X_ neurons display 2-4 rebound bursts that vary in their strength and duration due to HVC_INT_ – HVC_X_ inhibitory coupling as well as intrinsic properties (Fig. 9).

**Figure 9.**
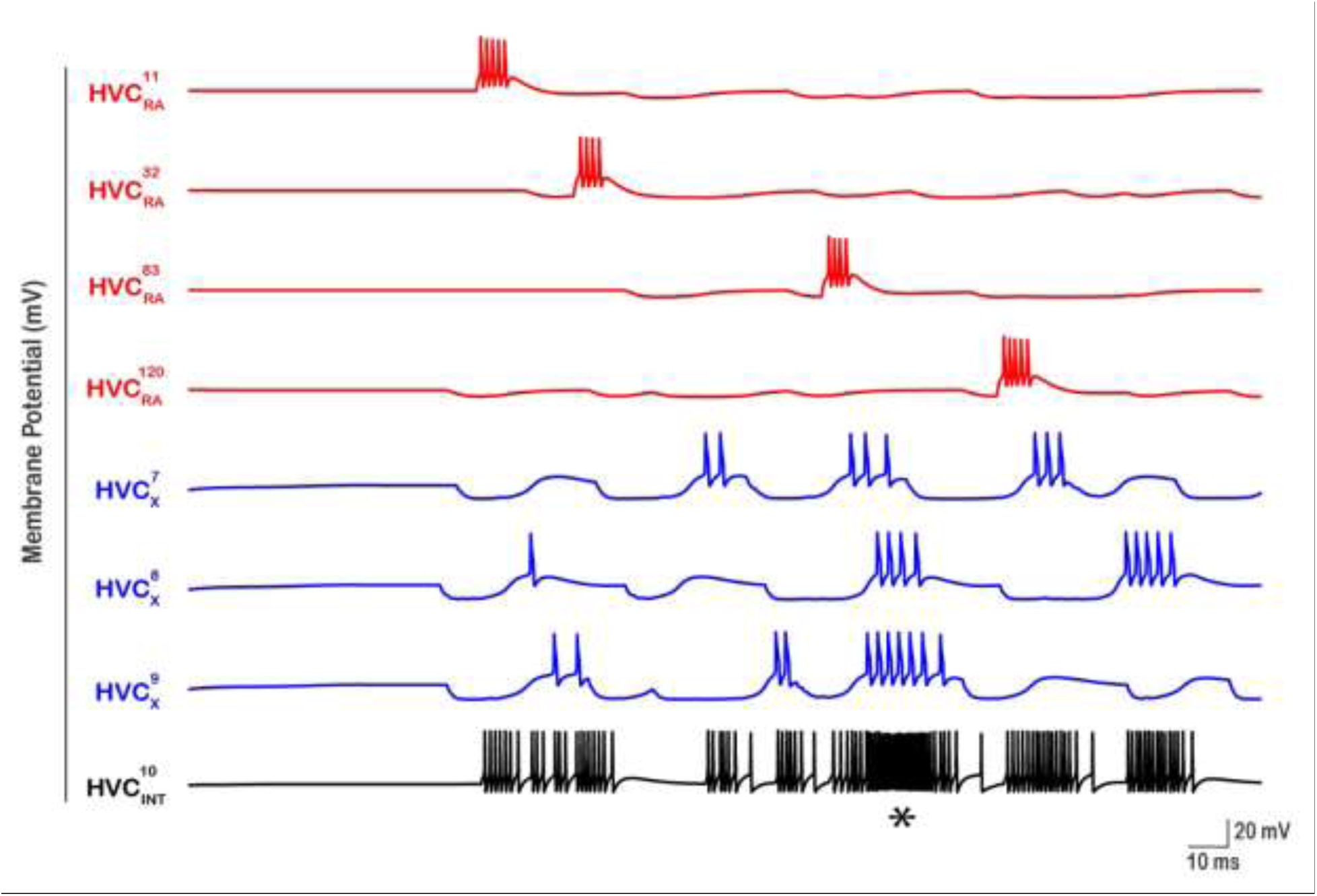
Patterned activity of HVC interneurons illustrated for an exemplar interneuron 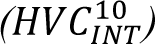. For this neuron, 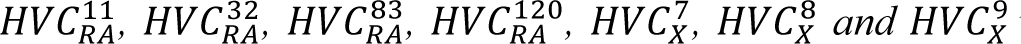 were selected randomly from the pool of HVC_RA_’s and HVC_X_’s to form excitatory coupling. The number of bursts in 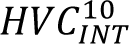 is controlled by the number of bursts that each of the HVC_RA_ and HVC_X_ neurons that connect to it exhibit. The strength of each of the 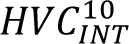 bursts depend on the magnitude of 𝑔_𝐴𝑀𝑃𝐴_ from the corresponding neuron(s) they cause it as well as the simultaneous bursting of any of the projecting neurons. For example, the asterisk (*) shows a region of dense firing in 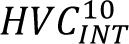 because 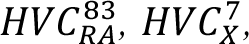, 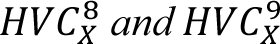 neurons elicit their spikes at similar times causing a potentiated response in 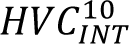. HVC_X_ neurons exhibit multiple sags and rebounds because they’re receiving inhibition from several interneurons (not shown here).

### Dense bursting in HVC_INT_

The random connections from HVC_RA_ and HVC_X_ neurons to HVC_INT_ as well as the multiple bursts HVC_INT_ and HVC_X_ exhibit in this network are illustrated in Figure 9 by highlighting a sample interneuron 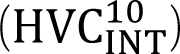. Here the firing patterns of 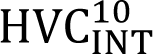 in addition to all HVC_RA_ and HVC_X_ neurons that connect to it are shown. 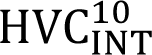 received random synaptic inputs from 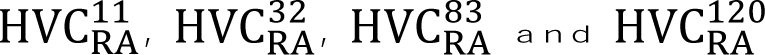 neurons, as well as from X-projecting neurons 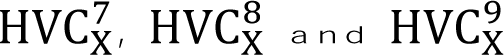 in the same microcircuit it belongs to. Notice that each HVC_RA_ neuron (which happen to belong to separate microcircuits in this case), bursts only once as reported experimentally (Hahnloser et al., 2002). 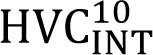 generates multiple bursts as well as spike sparsely. Each of the 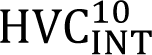 bursts are aligned with one of the bursts in HVC_RA_ and/or HVC_X_ neurons due to excitatory coupling.

The number of bursts an HVC_INT_ neuron exhibits is largely determined by the number of bursts that each HVC_RA_ and each HVC_X_ neuron that synapses onto that interneuron exhibits. In other words, the larger the number of bursts that a population of HVC_RA_ and HVC_X_ that connects to an interneuron exhibit, the larger the number of bursts elicited in the interneuron. The strength of the bursts in HVC_INT_ neurons is determined by two factors: 1) the strength of the excitatory synaptic conductance from HVC_RA_ and/or HVC_X_ to HVC_INT_, with stronger bursts in HVC_INT_ corresponding to larger magnitudes of g_AMPA_ from HVC_RA_ and/or HVC_X_ onto the interneuron, 2) the number of spikes/bursts in HVC_RA_ and HVC_X_ that are aligned and occur simultaneously at the HVC_INT_ synapses. For example, the asterisk shown in Figure 9 under the 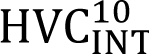 spike train highlights a stronger burst (compared to the rest of the spiking pattern) because 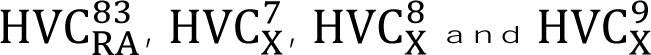 neurons tended to fire/burst at the same moment (or within a close interval of each other), thereby generating a stronger response in 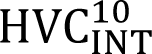. Therefore, the characteristic tonic activity that HVC interneurons exhibit with bursting and suppression at different locations during singing is explained in our model by the excitatory and inhibitory coupling between the interneurons and both classes of projecting neurons.

The inhibitory effect of HVC_INT_ neurons, their interplay with the excitatory projection neurons, and their intrinsic properties plays a key role in the modulation of sequence propagation and overall desired network behavior. In particular, if the excitation from the projection neurons onto the interneurons was very large (>> g_AMPA_), then HVC_INT_ neurons enter regimes of very dense and continuous bursting/spiking (above their natural and basic, yet already enhanced potentiation), thereby leading to the inhibition of the HVC_RA_’s, halting the sequence. Similarly, if the GABAergic conductances from HVC_INT_ to HVC_RA_ was relatively strong, outside their ideal ranges (Figure 2-B), then HVC_RA_ neurons won’t be able to elicit their bursts.

Besides the synaptic modulations of HVC_INT_ neurons on network activity, intrinsic mechanisms orchestrated primarily by the T-type Ca^2+^ current and the hyperpolarization-activated inward current need to be tightly regulated to ensure desired network activity. The T-type calcium channel opens near resting membrane potential and markedly influence neuronal excitability (Huguenard, 1996; Jagodic et al., 2008). In our model, if the T-type Ca^2+^ current in an HVC_INT_ neuron is up-regulated (due to a depolarizing shift in its voltage-dependent inactivation or simply setting g_CaT_ to a relatively large value), then the interneuron will fire with much larger firing frequency, silencing the corresponding HVC_RA_ and HVC_X_ neurons that it connects to and breaking the sequence (Fig. 10A). Similarly, the H-channels regulate the resting membrane potentials of the neurons they’re expressed in and play a key role in regulating the spontaneous firing activity (Datunashvili et al., 2018; Funahashi et al., 2003; Yao et al., 2003). In our model, if the H conductance of an HVC_INT_ was upregulated (by increasing the magnitude of g_h_), then the neuron switches to continuous firing, silencing the HVC_RA_’s that is connects to (Fig. 10B). Therefore, both the interplay between excitation and inhibition between HVC_INT_ and HVC projection neurons as well as the intrinsic properties of HVC_INT_ neurons (I_CaT_ and I_H_) are necessary elements to ensure an accurate propagation of sequential activity.

**Figure 10.**
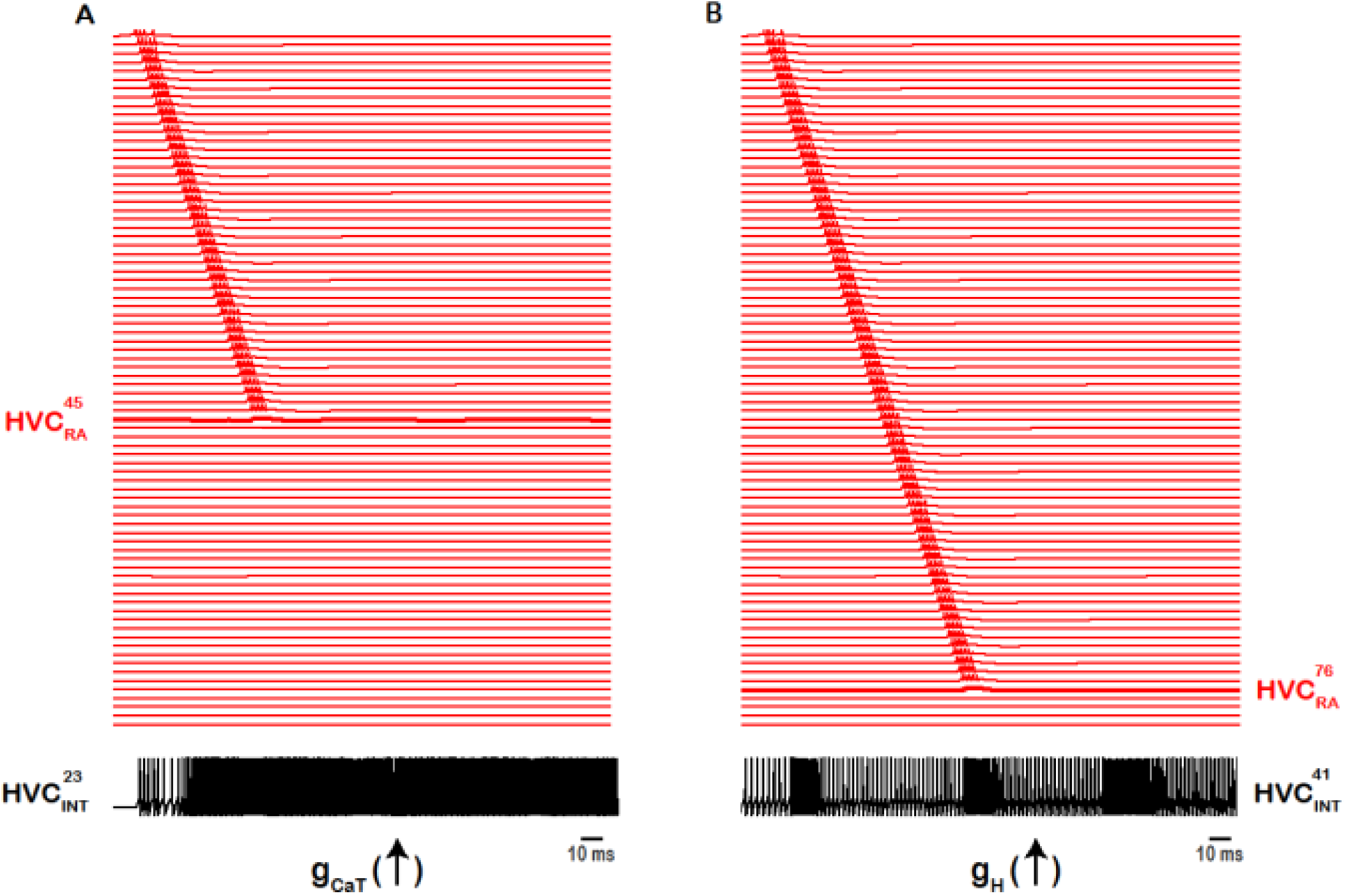
Intrinsic changes in HVC_INT_ halts the propagation of sequential activity. Up-regulating the T-type Ca^2+^ current conductance (**A**) or the hyperpolarization-activated inward current conductance (**B**) in exemplar HVC_INT_ neurons eliminates sequence propagation. Increasing 𝑔_C𝑎𝑇_ in 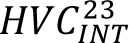 10-fold results in dense bursting and firing in 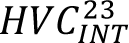, which in its turn blocks the bursting of 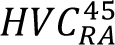 due to the inhibitory GABA coupling between 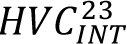 and 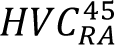 (**A**). Similarly, increasing 𝑔_𝐻_ in 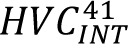 10-fold results in increased firing in 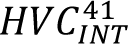, which in its turn blocks the bursting of 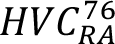 due to the inhibitory GABA coupling between 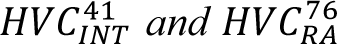 (**B**). Sequence of HVC_RA_ bursts truncated at the level of 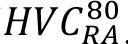 for better visualization purposes.

### Rebound bursting in HVC_X_

The activity patterns that HVC_X_ neurons exhibit in the network are illustrated in Figure 8B (shown here for 10 HVC_X_ neurons). The X-projecting neuronal firing is characterized by regions of inhibition throughout singing, interrupted by occasional rebound bursting. The strength of each burst and the number of total bursts in an HVC_X_ depend on synaptic and intrinsic mechanisms summarized briefly as such: 1) the degree of inhibition from HVC_INT_ neurons, 2) the number and timing of bursts of HVC_INT_ neurons and 3) the intrinsic properties and magnitudes of the T - type Ca^2+^ and H – currents that the HVC_X_ neurons exhibit. In short, the stronger the GABAergic maximal conductance(s) from the HVC_INT_ neurons that inhibit a given HVC_X_ neuron, the stronger the HVC_X_ neuron’s rebound burst (Figures 8-11). Similarly, if multiple HVC_INT_ neurons’ bursts that inhibit an HVC_X_ neuron aligned simultaneously, then the rebound in HVC_X_ is potentiated due to the stronger inhibition. This is primarily due to the T-type Ca^2+^ current and the H-current that HVC_X_ neurons exhibit, facilitating rapid calcium influx into the neurons when they rebound from hyperpolarization. The calcium influx is correlated to the degree and the duration of inhibition that the neuron receives, and can trigger a potentiated burst of action potentials leading to more robust rebound responses. In other words, the stronger the inhibition of HVC_X_, the stronger the activation of I_CaT_ and I_h_, leading to stronger rebounds. Moreover, the number of spikes in each HVC_X_ burst is controlled by several factors including 1) the degree of inhibition from HVC_INT_ and its effect on I_CaT_ and I_h_ as described earlier, and 2) the strength of the Ca^2+^-dependent K^+^ conductance (g_SK_) with stronger magnitudes of this conductance dampening the excitability of these neurons and reducing the number of spikes.

The interplay between HVC_INT_ and HVC_X_ in shaping the characteristic HVC_X_ responses is illustrated in Figure 11 from exemplar neurons in the network. Two interneurons, 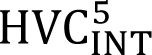 (black) and 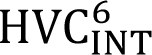 (green) were selected randomly to inhibit 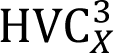 (blue trace, Fig. 11A). There are several regions in the 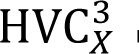 membrane potential trajectory where the neuron is inhibited (sags/sinks in the voltage trace). These regions correspond to the moments in time when either 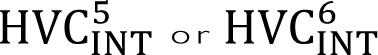 neurons, or both simultaneously, are bursting, thereby silencing 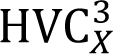. Eventually, 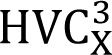 is able to escape inhibition at a few instances in time and generate multiple post inhibitory rebound bursts mediated by I_CaT_ and I_h_. Figure 11B shows a zoomed version of panel A illustrating the three rebound bursts in 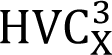 as a result of escape from inhibition. 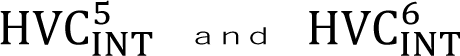 neurons generated multiple successive bursts of firing, during which 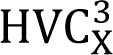 cannot escape the inhibition. There are only a few intervals where 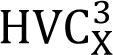 is able to elicit rebound bursts; the first opportunity was at the beginning of the trace (Fig. 11A) where the sag generated in 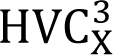 as a result of inhibition from 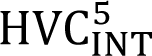 is clear, but the inhibition from 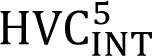 was not strong enough to elicit a rebound burst in 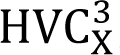, rather eliciting only a rebound depolarization. The subsequent three opportunities where 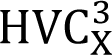 is able to escape the inhibition all elicited rebound bursts due to the strong inhibitory input arriving from both interneurons.

**Figure 11.**
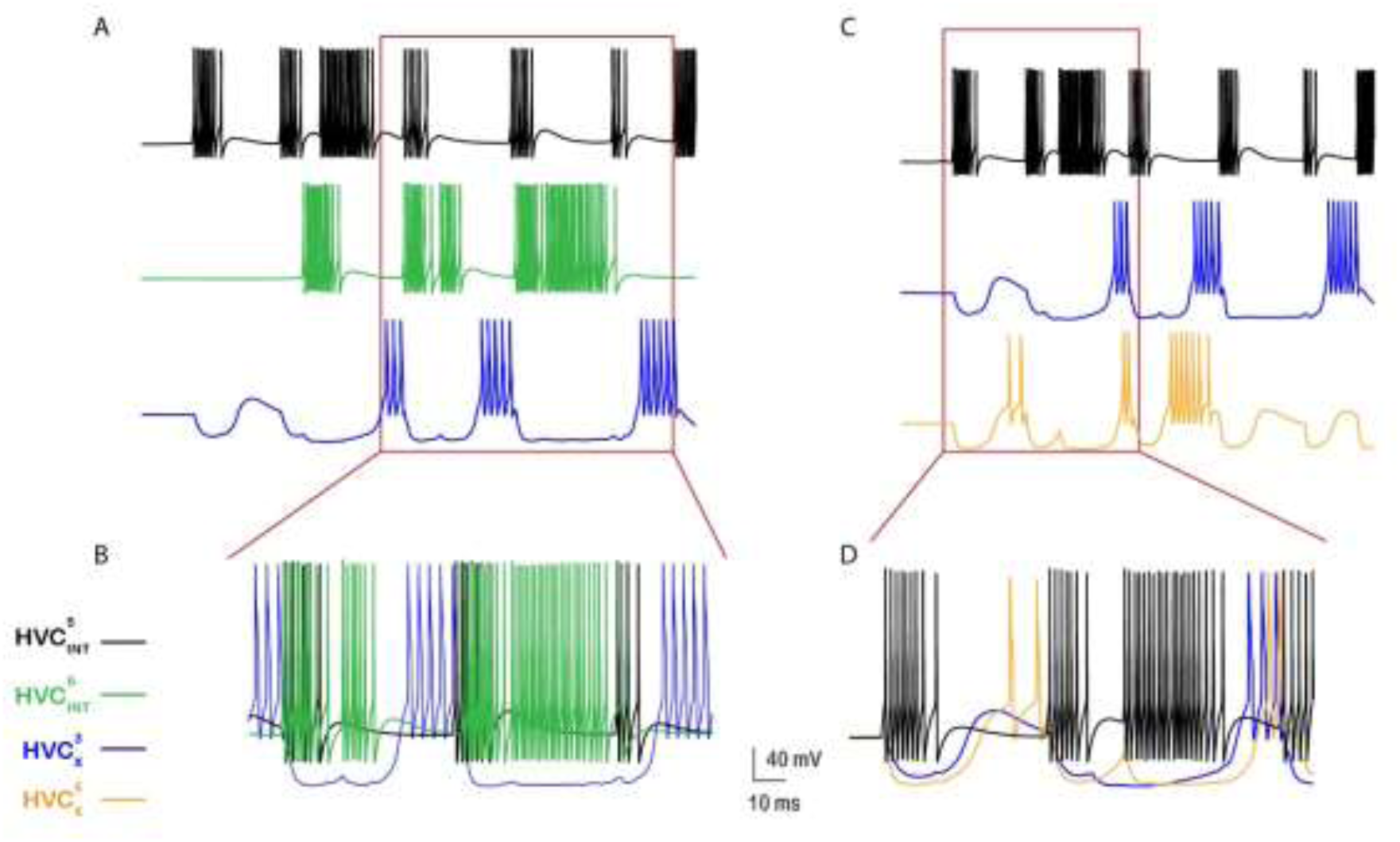
Activity patterns illustrating the interplay between HVC interneurons and X-projecting neurons. 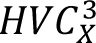 (blue) is an exemplar projecting neuron that receives inhibition from 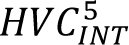 (black) and 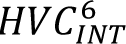 (green) due to the random inhibitory coupling (**A**). 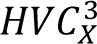 is inhibited whenever 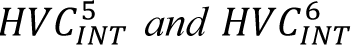 are firing, eventually escaping inhibition at some intervals and eliciting rebound bursts due to the activation of I_𝐻_ and I_C𝑎𝑇_ . HVC interneurons can inhibit multiple HVC_X_ neurons. 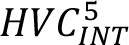 is an exemplar from the network that inhibits 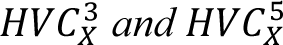 (orange) (**C**). Bursts in 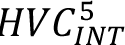 elicit subsequent bursts in 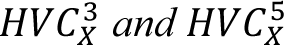 unless silenced by other HVC_INT_ neurons that connect to them. Zoomed versions of (**A**) and (**C**) are show in (**B**) and (**D**).

All rebound bursts in HVC_X_ are orchestrated by the hyperpolarization-activated inward current (I_H_) and the T-type Ca^2+^ current (I_CaT_). The longer the duration of the inhibition prior to a rebound, the longer the time I_H_ and I_CaT_ take to fully activate and generate the rebound burst. The shorter the inhibitory silent interval (that is, the interval in which HVC_X_ is receiving no inhibition), the shorter the rebound and the fewer the number of spikes in the rebound burst (Fig. 11A, first rebound burst). Moreover, HVC interneurons inhibit multiple HVC_X_ neurons in our network. Figure 11C shows an exemplar interneuron (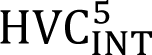, black) that happens to inhibit two X-projecting neurons (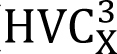, blue and 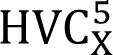, orange). Each burst in 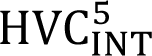 elicits an iPSP in 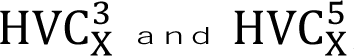, and can contribute to subsequent rebound bursts in them unless silenced by other interneurons that connect to 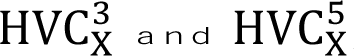 (Fig. 11D). All HVC_X_ neurons differed in their corresponding number of rebound bursts as well as in the number of spikes per burst rendering the results biologically accurate.

As mentioned earlier, the T-type Ca^2+^ and the H-conductances play a significant role in modulating the rebound bursting in HVC_X_ neurons. Due to this interplay, we do not need significant inhibition to generate rebound bursts, because the T-type Ca^2+^ current’s conductance can be stronger than the inhibitory conductance, leading to robust rebound bursting, even when the degree of inhibition is not very strong. As a matter of fact, these two conductances, along with the Ca^2+^ - dependent K^+^ conductance can halt the sequential propagation of activity and mess up the overall desired network behavior as described next. Ramping up g_H_ in HVC_X_ neurons to values outside the allowed ranges (Figure 2-A) will break the sequence of propagation and generate nonrealistic firing patterns. The g_H_ parameter was increased in Figure 12A in an exemplar HVC_X_ neuron 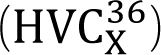 to large values, driving 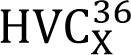 into regimes of runaway excitation due to the up-regulation of I_H_ and generating a non-realistic firing pattern for a typical X-projecting neuron during singing. The reason the sequential propagation of HVC_RA_ is halted is because 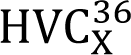 happens to have an excitatory coupling with 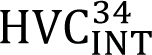 (along with other interneurons, but we illustrate it here for 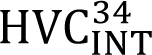), and as a result of 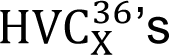 increased firing, 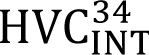 generates a mostly-continuous firing trace of bursting and spiking, which silences the HVC_RA_ neuron that it happens to inhibit (in this case, 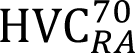, Fig. 12A).

**Figure 12.**
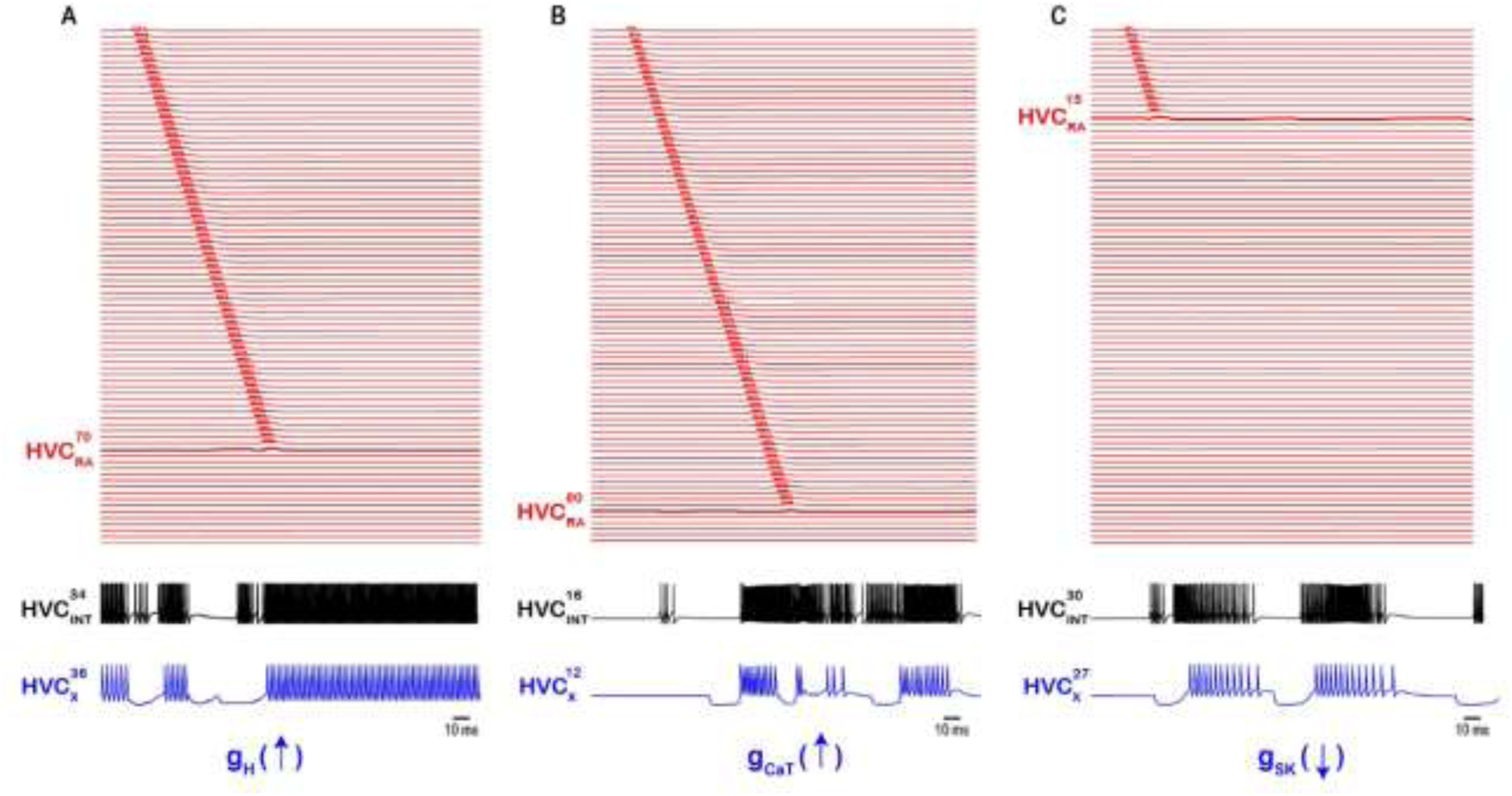
Intrinsic changes in HVC_X_ halts the propagation of sequential activity. **A**. Up-regulating the hyperpolarization-activated inward current conductance in a sample HVC_X_ neuron (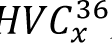, by increasing its 𝑔_H_ 10-fold) leads to increased firing in all HVC_INT_ neurons it connects to (for example, 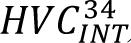), which in its turn inhibits all HVC_RA_ neurons it connects to (for example, 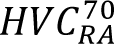, being first in the pool that it inhibits) breaking the sequence at the level of 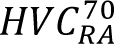. **B**. Up-regulating the T-type Ca^2+^ current conductance in a sample HVC_X_ neuron (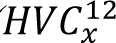, by increasing its 𝑔_C𝑎𝑇_ 15-fold) leads to stronger rebound bursts in 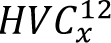 which leads to increased firing in all HVC_INT_ neurons it connects to (for example, 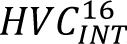), which in its turn inhibits all HVC_RA_ neurons it connects to (for example, 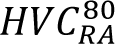, being first in the pool that it inhibits) breaking the sequence at the level of 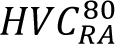. **C**. Finally, down-regulating the Ca^2+^ - dependent K^+^ current conductance in a sample HVC_X_ neuron (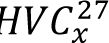, by setting its 𝑔_SK_ to zero) leads to stronger rebound bursts in 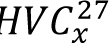 which leads to increased firing in all HVC_INT_ neurons it connects to (for example, 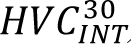), which in its turn inhibits all HVC_RA_ neurons it connects to (for example, 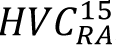, being first in the pool that it inhibits) breaking the sequence at the level of 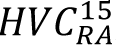. Sequence of HVC_RA_ bursts truncated at the level of 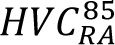 for better visualization purposes.

Similarly, altering g_CaT_ in HVC_X_ neurons can have similar effects on network activity. Figure 12B shows an exemplar HVC_X_ neuron (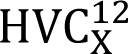) where g_CaT_ was ramped up to large values and as a result, stronger rebound bursts in 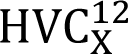 were elicited as well as larger number of rebound spikes in each burst. This is primarily due to the up-regulation of the T-type Ca^2+^ current that was induced, markedly influencing the neuronal excitability. The stronger rebounding in 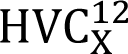 generated stronger excitation in their postsynaptic interneurons’ counterparts that they excite (for example, 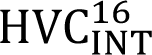), which in their turn silenced the HVC_RA_ neurons that they inhibit (for example, 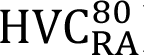 is the first neuron in the chain that 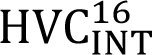 inhibits, breaking the sequence at 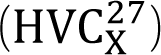). And finally, the Ca^2+^ - dependent K^+^ conductance, which plays a key role in governing HVC_X_ neurons’ excitability and their characteristic spike frequency adaptation (Daou et al., 2013) can have similar effects if the channel was down-regulated (rather than up-regulated as in I_CaT_ and I). Figure 12C shows the effects of blocking the g_SK_ conductance in an exemplar HVC_X_ neuron (HVC^27^) leading to its increased firing, which in turn leads to stronger bursting and spiking in the HVC_INT_ neurons it excites (for example, 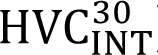). 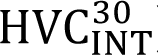 in its turn generates stronger and wider-range inhibition onto the HVC_RA_ neurons it sends its axons to (for example, 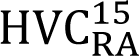 is the first neuron in the chain 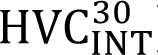 inhibits, silencing it and breaking the sequence at 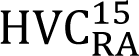).

Finally, we checked the consequences of altering the intrinsic properties of HVC_X_ neurons on the network’s desired behavior. To do so, we varied the maximal conductances of the three principal ionic currents of the X-projecting neurons (I_CaT_, I_SK_, I_H_) across all neurons of the population, while keeping this variation within the reported ranges shown in Figure 2A (because certainly going outside these ranges will disrupt network activity for other reasons as reported in Figures 10 and 12). Varying those three key paramters across the HVCx population had different results. In 100 different simulations that generated random maximal conductances for I_CaT_,I_SK_ and I_H_, 41% of the simulations did not have any considerable effect on the desired network activity, whereas 59% resulted in disrupted network activity, and sometimes detrimental. For example, Figure 13 shows an example where the sequential propagation of activity was halted and the firing patterns of some interneurons and X-projecting neurons were rendered non-biophysically realistic. Particularly, some interneurons switched to continuous spiking or phasic bursting with little episodic bursting modes, while some HVC_X_ neurons generated longer rebounds, fewer number of bursts or fewer number of spikes per burst (Fig. 13 B-C). Hence, changes in the intrinsic properties of X-projecting neurons can disrupt activity propagation necessary for song production, and produce biologically unrealistic bursting patterns in HVC neurons. This can wreak havoc on our network model hinting to the finding that biophysical parameters are distinct and consistent for an individual bird and this unique combination is needed for song (Daou & Margoliash, 2020). The homogeneity in the intrinsic properties of X-projectors might be a strategy allowing it to adapt or respond to changes in the network.

**Figure 13.**
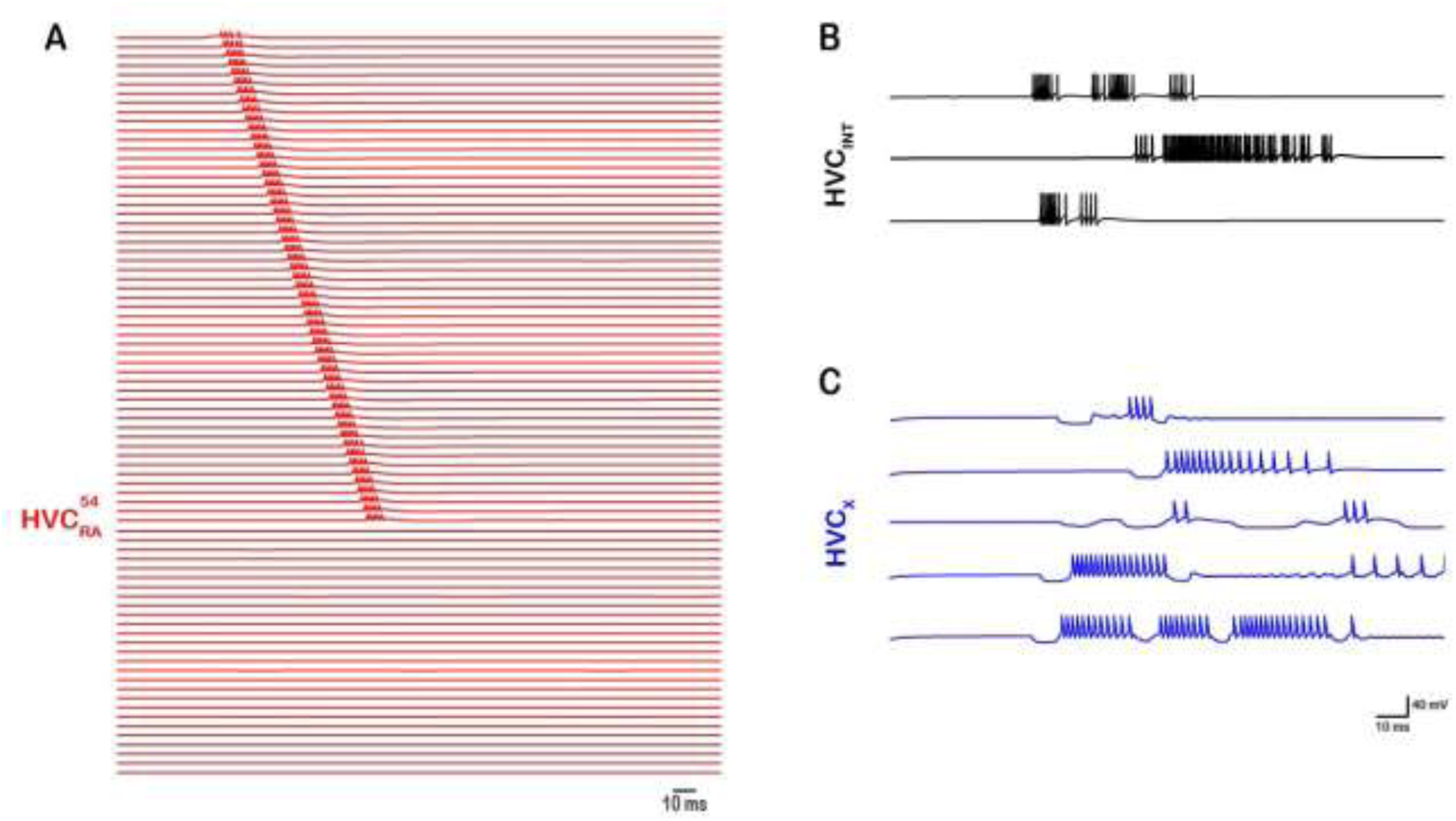
Altering the intrinsic properties of HVCx neurons disrupt network activity in 59% of the cases (out of 100 simulations) where the maximal conductances (gCaT, gSK and gH) of HVCx neurons are randomly varied within their allowed ranges (Figure 2A). As a result, some HVC interneurons (B) and X-projecting neurons (C) generated non-biological realistic firing patterns, halting the propagation of sequential activity in RA-projectors (A).

In conclusion, we developed a detailed and biophysically realistic neural network model for sequence propagation in the HVC of the zebra finch. Our model consisted of chains of microcircuits, each comprised of a selection of HVC_RA_, HVC_INT_ and HVC_X_ model neurons selected randomly from a total pool of neurons. The maximal conductances of the four key and principle ionic currents for each model neuron, the number of neurons of each class in any microcircuit, and the excitatory and inhibitory connections between the different classes within a microcircuit and across microcircuits are all selected randomly. This activity propagates throughout the chain of microcircuits causing a sequence of HVC_RA_ bursts while leaving behind realistic bursting patterns for all classes of HVC neurons as seen during singing. The model incorporates all known ionic and synaptic currents for each HVC neuron. The network architecture we developed was able to replicate the *in vivo* biologically realistic firing behavior for each class by including sparse timely-locked bursting in the RA-projecting neurons (with accurate intrinsic properties for each burst in terms of number of spikes, duration and spike morphology), multiple bursting in the X-projecting neurons that are also sparse and time-locked, and dense bursting/spiking in the interneurons with few intermittent quiescence. The ability of our network to reproduce the sequential propagation of activity in the presence of excitatory and inhibitory connections involving all neuronal subclasses as well as over a range of values for each synaptic and ionic maximal conductances is an indication of its robustness. Our network unveiled key intrinsic and synaptic mechanisms that modulate the sequential propagation of neural activity by highlighting important roles for the T-type Ca^2+^ current and hyperpolarization activated (H) inward current in HVC_X_ and HVC_INT_ neurons, Ca^2+^-dependent K^+^ current in HVC_X_ and HVC_RA_, A-type K^+^ current in HVC_RA_, as well as GABAergic and glutamatergic synaptic currents that connects all neuronal subclasses together. The result is an improved characterization of the HVC network responsible for song production in the zebra finch. Beyond replicating established HVC firing patterns, our model provides testable hypotheses that intrinsic membrane properties, particularly inhibitory timing and rebound bursts maintain robust sequential propagation. This context generates clear experimental predictions: for example, modulating I_H_ or I_CaT_ in HVC_X_ or HVC_INT_ neurons directly affects rebound spiking and sequence propagation; or altering I_SK_ or I_A_ conductances disrupts the ability of HVC_RA_ neurons to burst at the right time. The model also suggests that structured inhibition can act as a temporal scaffold for burst timing guiding experiments that manipulate interneuron dynamics or the overall network inhibition through optogenetics or pharmacology. Hence, a significant strength of the proposed model is that it outs forward suggestions for experimental manipulations in the form of targeted experiments whether using optogenetics or pharmacology that would help validate the model’s mechanisms and further clarify the specific roles different HVC components play in driving sequential activity.

## Discussion

In this study, we developed a biophysically realistic neural network model to explore how intrinsic neuronal properties and local connectivity within the songbird nucleus HVC may support the generation of temporally precise activity sequences associated with zebra finch song. The biophysically realistic network architecture that we designed combine both classes of HVC projection neurons with local inhibitory interneurons. A fundamental goal that we have achieved in our design is a successful replication of the *in vivo* firing behaviors of all the HVC neuronal classes: single sparse timely-precise bursting (3-6 spikes for ∼10 ms) in the RA-projecting neurons, multiple bursting (1-4 bursts with 4-9 spikes/burst) in the X-projecting neurons, dense and frequent bursting in the interneurons, as well as the general intrinsic properties that each class of HVC neurons exhibit (Daou et al., 2013; Lewicki, 1996; Long et al., 2010). The patterning activity in HVC is largely shaped in our model by the intrinsic properties of the individual neurons as well as the synaptic properties where excitation and inhibition play a major role in enabling neurons to generate their characteristic bursts during singing.

The three classes of model neurons incorporated to our network as well as the synaptic currents that connect them are based on Hodgkin-Huxley formalisms that contain ion channels and synaptic currents which had been pharmacologically identified (Daou et al., 2013; Kosche et al., 2015; Mooney & Prather, 2005). Our network showed that sequence propagation can be broken if several intrinsic mechanisms are perturbed. In particular, if I_CaT_ or I_H_ are upregulated in HVC_X_ or HVC_INT_, if I_SK_ is downregulated in HVC_X_ or if I_SK_ is upregulated in HVC_RA_, then the corresponding chain of activity stops and the rhythmic activity of the network is disrupted (Figures 7, 10 and 12). Synaptically, perhaps the most critical role in our network design is played by interneurons which orchestrate the activity of the two projection neurons in a structured manner. Interneurons adjust the timing of HVC projection neurons’ bursts (Amador et al., 2013; Kosche et al., 2015), and developmental learning regulates inhibition onto HVC_RA_ (Vallentin et al., 2016). While findings of Kosche et al. (2015) emphasize the robustness of the HVC timing circuit to inhibition, our model is more sensitive to inhibition, highlighting that HVC likely operates with several, redundant mechanisms that overall ensure temporal precision. HVC_RA_ neurons interact with HVC_X_ through local interneurons, a disynaptic inhibitory pathway that conveys information to HVC_X_ neurons (Prather et al., 2008). Focal application of the GABA_A_ receptor antagonist, gabazine, restricted inhibitory impact in HVC leading to stronger and faster responses relative to call onset (Benichov & Vallentin, 2020), showing that local HVC interneurons form an inhibitory mask that can greatly constrain the spiking activity of projecting neurons (Kornfeld et al., 2017; Kosche et al., 2015; Mooney & Prather, 2005) suggesting that HVC model networks that lack inhibitory neurons are inadequate for explaining sequential propagation of neural activity.

Various models of how the song is encoded within HVC have been proposed. Some groups suggested that bursting activity propagates through a chain of synaptically connected HVC_RA_ neurons either as single neurons (Fee et al., 2004; Hahnloser et al., 2002; Long et al., 2010) or as pools of HVC_RA_ neurons, each group driving a distinct ensemble of RA neurons (Jin et al., 2007; Leonardo & Fee, 2005; Li & Greenside, 2006).

These models assume that HVC_RA_ neurons generate a continuous, feed-forward sequence of activity over time, with little or no role played by X-projecting HVC neurons and interneurons. Other models have incorporated alternative temporal encoding mechanisms by necessitating synaptic integration at the levels of HVC_RA_ and HVC_INT_ populations (Drew & Abbott, 2003; Gibb et al., 2009a; Jin, 2009; Weber & Hahnloser, 2007) while yet other approaches gave emphasis to brainstem feedback processes by incorporating inter-hemispheric coordination to activate sequences of syllable-specific HVC_RA_ and HVC_INT_ neurons (Galvis et al., 2018; Gibb et al., 2009b). A prominent model used spatially recurrent excitatory chains and local feedback inhibition to show how the HVC network stabilize synchrony while propagating sequential activity (Cannon et al., 2015; Markowitz et al., 2015).

All existing models that describe premotor sequence generation in the HVC either assume a distributed model (Elmaleh et al., 2021) that dictates that local HVC circuitry is not sufficient to advance the sequence but rather depends upon moment-to-moment feedback through Uva (Hamaguchi et al., 2016), or assume models that rely on intrinsic connections within HVC to propagate sequential activity. In the latter case, some models assume that HVC is composed of multiple discrete subnetworks that encode individual song elements (Glaze & Troyer, 2013; Long & Fee, 2008; Wang et al., 2008), but lacks the local connectivity to link the subnetworks, while other models assume that HVC may have sufficient information in its intrinsic connections to form a single continuous network sequence (Long et al., 2010).

The network architecture we developed here exhibits overlap with the various models presented. First, in agreement with the continuous model, our network architecture displays a feed-forward mechanism regulating the circuit dynamics e.g., (Gibb et al., 2009a; Jin, 2009; Jin et al., 2007; Li & Greenside, 2006; Long et al., 2010). Nonetheless, diverging from a linear progression of HVC neurons directing the song, the network’s structure comprises sequences of microcircuits incorporating all classes of HVC neurons, where sequential activity transmits from one microcircuit to the next, as opposed to transitioning directly between individual neurons. Second, in agreement with the subnetwork models, our model envisions HVC as comprised of multiple discrete subcircuits (SSSs) where each microcircuit incorporates its own pool of neurons; however, in our model HVC’s connectivity is sufficient to link the microcircuits together and extrinsic influences are not needed. Moreover, our network is in agreement with the (Cannon et al., 2015) model where structured inhibition is needed to propagate sequential activity, synchronize the firing of pools of neurons and stabilize spike timing along the chain. The pivotal element in advancing sequential activity through time is the inhibition exerted by HVC_INT_ onto HVC_X_ and HVC_RA_ neurons, facilitated in the case of HVC_X_ through rebound firing, and all orchestrated by intrinsic mechanisms.

A potential drawback of our model is that it does not incorporate brainstem feedback processes or address inter-hemispheric coordination as proposed by others (Galvis et al., 2018; Gibb et al., 2009b). Another drawback is its sole focus on local excitatory connectivity within the HVC (Kornfeld et al., 2017; Long et al., 2010). Moreover, HVC neurons receive afferent excitatory connections (Akutagawa & Konishi, 2010; Nottebohm et al., 1982) that plays significant roles in their local dynamics. For example, the excitatory inputs that HVC neurons receive from Uvaeformis may be crucial in initiating (Andalman et al., 2011; Danish et al., 2017; Galvis et al., 2018) or sustaining (Hamaguchi et al., 2016) the sequential activity. In addition, while our simplified, somatically driven architecture enables better exploration of mechanisms for sequence propagation, future extensions of the model will incorporate dendritic compartments to more accurately reflect the intrinsic bursting mechanisms observed in HVC_RA_ neurons. Moreover, our model was run at a fixed physiological temperature, but it’s well known going all the way back to Hodgkin and Huxley that both ion channel activity and synaptic dynamics can change with temperature. In future work, adding temperature scaling (like Q10 factors) could help us explore how burst timing and sequence speed change with temperature changes, and how neural activity in HVC would/would not preserve its precision under different physiological conditions.

### The role of ion channels in controlling network activity

Our model highlights the role of principal ion channels (I_H_, I_CaT_, I_SK_ and I_A_) in controlling HVC’s network dynamics and progressing its neural sequence. Hyperpolarization-activated ionic conductances had been widely observed across various electrically excitable cells (Pape, 1996) and play significant roles in rhythmogenesis (Budde et al., 1997; Golowasch et al., 1992; Golowasch & Marder, 1992). In our network, model HVC_X_ neurons are not able to elicit their rebound bursting without I_H_ and sequence is halted if this conductance is upregulated in either of HVC_X_ or HVC_INT_ (Figures 8-10, 11 and 12). Similarly, the T-type Ca^2^+ current is recognized as crucial in various systems as an ionic contributor to burst generation (Deschenes et al., 1982; Fraser & MacVicar, 1991; Huguenard, 1996; Llinás & Yarom, 1981). Lewicki (1996) observed a significant hyperpolarization in some HVC neurons *in vivo* before they emit their corresponding bursts, and the intensity of the burst correlates with the degree of hyperpolarization. In this study, we have illustrated its pivotal role in rebound spiking where up-regulating this conductance in HVC_X_ or HVC_INT_ halts sequence propagation (Figures 8-10, 11 and 12). Moreover, the A-type K^+^ current is involved in several rhythmogenic activities controlling membrane excitability (Coetzee et al., 1999; Ellis et al., 2007; Gross et al., 2016) and in our network, upregulating I_A_ suppress bursting in model HVC_RA_ and breaks sequence propagation (Fig. 7). Finally, the small conductance Ca^2+^-activated potassium current (I_SK_) plays important roles in the regulation of excitable cells controlling network rhythmic activity (Benítez et al., 2011; Chen et al., 2014; Pedarzani et al., 2005) and in our network I_SK_ plays a significant role since its upregulation in HVC_RA_ or its downregulation in HVC_X_ eliminates sequence propagation (Fig.7 and Fig. 12 respectively).

In conclusion, the network model developed provide a large step forward in describing the biophysics of HVC circuitry, and may throw a new light on certain dynamics in the mammalian brain, particularly the motor cortex (Shmiel et al., 2006) and the hippocampus regions (Lee & Wilson, 2002) where precisely-timed sequential activity is crucial. We suggest that temporally-precise sequential activity may be a manifestation of neural networks comprised of chain of microcircuits, each containing pools of excitatory and inhibitory neurons, with local interplay among neurons of the same microcircuit and global interplays across the various microcircuits, and with structured inhibition and intrinsic properties synchronizing the neuronal pools and stabilizing timing within the ongoing sequence.

## APPENDIX 1

### Voltage-gated ionic currents

The constant-conductance leak current is *I*_𝐿_ = 𝑔_𝐿_(𝑉−𝑉_𝐿_). The remaining voltage-gated ionic currents have non-constant dependent currents with activation/inactivation kinetics:

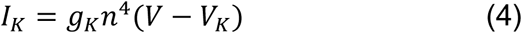

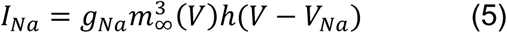

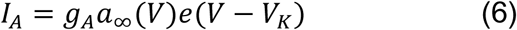

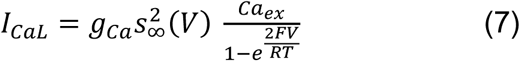

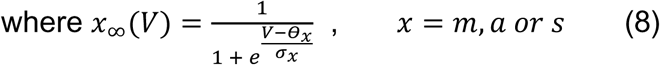

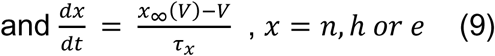

where 𝑥_∞_(𝑉) for n and e is given by (8) and for ℎ_∞_ as follows

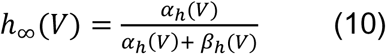

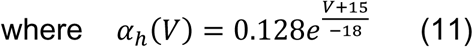

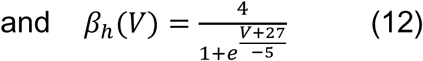

### Low-voltage activated T-type calcium current

The low-voltage activated T-type Ca2+ current is described by the Goldman-Hodgkin-Katz formula:

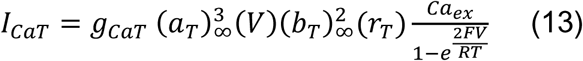

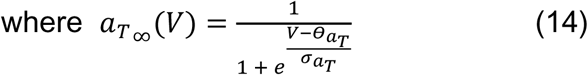

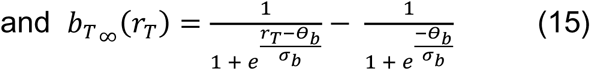

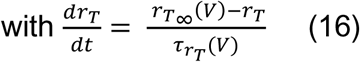

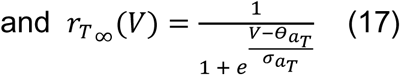

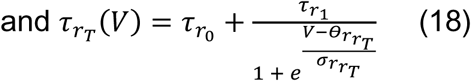

### Calcium-dependent Potassium current

The small conductance potassium current (*I*_𝑆𝐾_) is modeled as

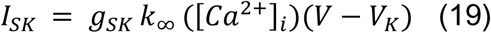

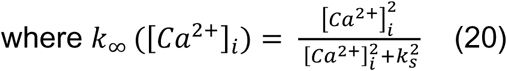

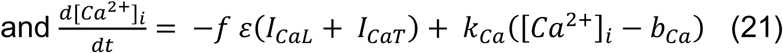

### Hyperpolarization-activated inward current

The hyperpolarization activated inward current’s activation is modeled as in Destexhe and Babloyantz (1993) using a fast component (𝑟_𝑓_) and a slow component (𝑟_𝑠_) as follows:

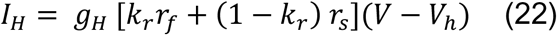

The fast activation component is given by:

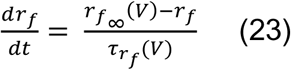

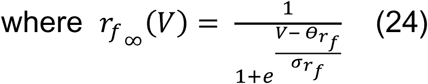

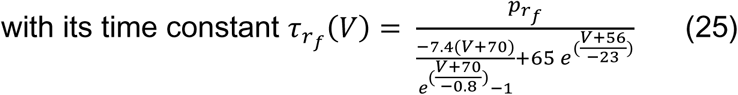

The slow activation component obeys:

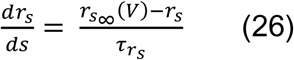

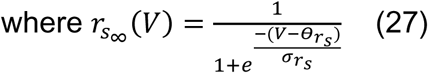

Synaptic currents equations

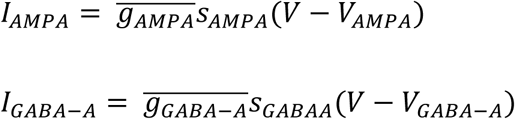

where 𝑠_𝐴𝑀𝑃𝐴_ and 𝑠_𝐺𝐴𝐵𝐴𝐴_ are given by

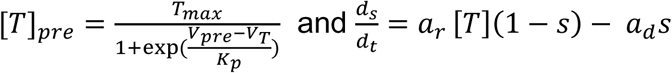

with 𝑇_*m*𝑎𝑥_ = 1, 𝐾_𝑝_ = 5, 𝑉_𝑇_ = 2. For GABA_A_, 𝑎_𝑟_ = 5 and 𝑎_𝑑_ = 0.18, while for AMPA, 𝑎_𝑟_ = 1.1 and 𝑎_𝑑_ = 0.19.

## APPENDIX 2

**Table 1.** Fixed parameter values used in all simulations.

| Parameter | Value | Parameter | Value |
| --- | --- | --- | --- |
| $V_L$ | $-70\text{ mV}$ | $\theta_{aT}$ | $-65\text{ mV}$ |
| $V_K$ | $-90\text{ mV}$ | $\theta_b$ | $0.4\text{ mV}$ |
| $V_{Na}$ | $50\text{ mV}$ | $\theta_{rT}$ | $-67\text{ mV}$ |
| $V_{Ca}$ | $85\text{ mV}$ | $\theta_{r_{rT}}$ | $68\text{ mV}$ |
| $V_H$ | $-30\text{ mV}$ | $\sigma_m$ | $-5\text{ mV}$ |
| $g_L$ | $2\text{ nS}$ | $\sigma_n$ | $-5\text{ mV}$ |
| $g_{Ca}$ | $19\text{ nS}$ | $\sigma_s$ | $-0.05\text{ mV}$ |
| $\tau_n$ | $10\text{ msec}$ | $\sigma_a$ | $-10\text{ mV}$ |
| $\tau_{hp}$ | $1000\text{ msec}$ | $\sigma_e$ | $5\text{ mV}$ |
| $\tau_e$ | $20\text{ msec}$ | $\sigma_{rf}$ | $5\text{ mV}$ |
| $\tau_h$ | $1\text{ msec}$ | $\sigma_{rs}$ | $25\text{ mV}$ |
| $\tau_{rs}$ | $1500\text{ msec}$ | $\sigma_{aT}$ | $-7.8\text{ mV}$ |
| $\tau_{r0}$ | $200\text{ msec}$ | $\sigma_b$ | $-0.1\text{ mV}$ |
| $\tau_{r1}$ | $87.5\text{ msec}$ | $\sigma_{rT}$ | $2\text{ mV}$ |
| $\theta_m$ | $-35\text{ mV}$ | $\sigma_{r_{rT}}$ | $2.2\text{ mV}$ |
| $\theta_n$ | $-30\text{ mV}$ | $f$ | $0.1$ |
| $\theta_s$ | $-20\text{ mV}$ | $\varepsilon$ | $0.0015\text{ pA}^{-1}\text{ microM msec}^{-1}$ |
| $\theta_a$ | $-20\text{ mV}$ | $k_{Ca}$ | $0.3\text{ msec}^{-1}$ |
| $\theta_e$ | $-60\text{ mV}$ | $b_{Ca}$ | $0.1\text{ micro M}$ |
| $\theta_{rf}$ | $-105\text{ mV}$ | $k_s$ | $0.5\text{ micro }M$ |
| $\theta_{rs}$ | $-105\text{ mV}$ | $p_{rf}$ | 100 |
| $g_{Na_{HVCRA}}$ | $300\text{ nS}$ | $g_{K_{HVCRA}}$ | $150\text{ nS}$ |
| $g_{Na_{HVCX}}$ | $450\text{ nS}$ | $g_{K_{HVCX}}$ | $80\text{ nS}$ |
| $g_{Na_{HVCINT}}$ | $800\text{ nS}$ | $g_{K_{HVCINT}}$ | $1200\text{ nS}$ |

## References

1. Akutagawa, E., & Konishi, M. (2010). New brain pathways found in the vocal control system of a songbird. Journal of Comparative Neurology, 518(15), 3086–3100.

2. Amador, A., Perl, Y. S., Mindlin, G. B., & Margoliash, D. (2013). Elemental gesture dynamics are encoded by song premotor cortical neurons. Nature, 495(7439), 59–64.

3. Andalman, A. S., Foerster, J. N., & Fee, M. S. (2011). Control of vocal and respiratory patterns in birdsong: dissection of forebrain and brainstem mechanisms using temperature. PLoS One, 6(9), e25461.

4. Benichov, J. I., & Vallentin, D. (2020). Inhibition within a premotor circuit controls the timing of vocal turn-taking in zebra finches. Nature communications, 11(1), 221.

5. Benítez, B. A., Belálcazar, H. M., Anastasía, A., Mamah, D. T., Zorumski, C. F., Mascó, D. H., Herrera, D. G., & De Erausquin, G. A. (2011). Functional reduction of SK3-mediated currents precedes AMPA-receptor-mediated excitotoxicity in dopaminergic neurons. Neuropharmacology, 60(7-8), 1176–1186.

6. Budde, T., Biella, G., Munsch, T., & Pape, H. C. (1997). Lack of Regulation by Intracellular Ca2+ of the Hyper Polarization-Activated Cation Current in Rat Thalamic Neurones. The Journal of physiology, 503(1), 79–85.

7. Cannon, J., Kopell, N., Gardner, T., & Markowitz, J. (2015). Neural sequence generation using spatiotemporal patterns of inhibition. PLoS computational biology, 11(11), e1004581.

8. Chen, L., Deltheil, T., Turle-Lorenzo, N., Liberge, M., Rosier, C., Watabe, I., Sreng, L., Amalric, M., & Mourre, C. (2014). SK channel blockade reverses cognitive and motor deficits induced by nigrostriatal dopamine lesions in rats. International Journal of Neuropsychopharmacology, 17(8), 1295–1306.

9. Coetzee, W. A., Amarillo, Y., Chiu, J., Chow, A., Lau, D., McCormack, T., Morena, H., Nadal, M. S., Ozaita, A., & Pountney, D. (1999). Molecular diversity of K+ channels. Annals of the New York Academy of Sciences, 868(1), 233–255.

10. Danish, H. H., Aronov, D., & Fee, M. S. (2017). Rhythmic syllable-related activity in a songbird motor thalamic nucleus necessary for learned vocalizations. PLoS One, 12(6), e0169568.

11. Daou, A., & Margoliash, D. (2020). Intrinsic neuronal properties represent song and error in zebra finch vocal learning. Nature communications, 11(1), 952.

12. Daou, A., Ross, M. T., Johnson, F., Hyson, R. L., & Bertram, R. (2013). Electrophysiological characterization and computational models of HVC neurons in the zebra finch. Journal of neurophysiology, 110(5), 1227–1245.

13. Datunashvili, M., Chaudhary, R., Zobeiri, M., Lüttjohann, A., Mergia, E., Baumann, A., Balfanz, S., Budde, B., Van Luijtelaar, G., & Pape, H.-C. (2018). Modulation of hyperpolarization-activated inward current and thalamic activity modes by different cyclic nucleotides. Frontiers in cellular neuroscience, 12, 369.

14. Deschenes, M., Roy, J., & Steriade, M. (1982). Thalamic bursting mechanism: an inward slow current revealed by membrane hyperpolarization. Brain research, 239(1), 289–293.

15. Destexhe, A., & Babloyantz, A. (1993). A model of the inward current Ih and its possible role in thalamocortical oscillations. Neuroreport, 4(2), 223–226.

16. Destexhe, A., Mainen, Z. F., & Sejnowski, T. J. (1994). Synthesis of models for excitable membranes, synaptic transmission and neuromodulation using a common kinetic formalism. Journal of computational neuroscience, 1, 195–230.

17. Drew, P. J., & Abbott, L. (2003). Model of song selectivity and sequence generation in area HVc of the songbird. Journal of neurophysiology, 89(5), 2697–2706.

18. Dunmyre, J. R., Del Negro, C. A., & Rubin, J. E. (2011). Interactions of persistent sodium and calcium-activated nonspecific cationic currents yield dynamically distinct bursting regimes in a model of respiratory neurons. Journal of computational neuroscience, 31, 305–328.

19. Dutar, P., Vu, H. M., & Perkel, D. J. (1998). Multiple cell types distinguished by physiological, pharmacological, and anatomic properties in nucleus HVc of the adult zebra finch. Journal of neurophysiology, 80(4), 1828–1838.

20. Egger, R., Tupikov, Y., Elmaleh, M., Katlowitz, K. A., Benezra, S. E., Picardo, M. A., Moll, F., Kornfeld, J., Jin, D. Z., & Long, M. A. (2020). Local axonal conduction shapes the spatiotemporal properties of neural sequences. Cell, 183(2), 537–548. e512.

21. Ellis, L. D., Krahe, R. d., Bourque, C. W., Dunn, R. J., & Chacron, M. J. (2007). Muscarinic receptors control frequency tuning through the downregulation of an A-type potassium current. Journal of neurophysiology, 98(3), 1526–1537.

22. Elmaleh, M., Kranz, D., Asensio, A. C., Moll, F. W., & Long, M. A. (2021). Sleep replay reveals premotor circuit structure for a skilled behavior. Neuron, 109(23), 3851–3861. e3854.

23. Fee, M. S., & Goldberg, J. H. (2011). A hypothesis for basal ganglia-dependent reinforcement learning in the songbird. Neuroscience, 198, 152–170.

24. Fee, M. S., Kozhevnikov, A. A., & Hahnloser, R. H. (2004). Neural mechanisms of vocal sequence generation in the songbird. Annals of the New York Academy of Sciences, 1016(1), 153–170.

25. Fee, M. S., & Scharff, C. (2010). The songbird as a model for the generation and learning of complex sequential behaviors. ILAR journal, 51(4), 362–377.

26. Fraser, D. D., & MacVicar, B. A. (1991). Low-threshold transient calcium current in rat hippocampal lacunosum-moleculare interneurons: kinetics and modulation by neurotransmitters. Journal of Neuroscience, 11(9), 2812–2820.

27. Fujimoto, H., Hasegawa, T., & Watanabe, D. (2011). Neural coding of syntactic structure in learned vocalizations in the songbird. Journal of Neuroscience, 31(27), 10023–10033.

28. Funahashi, M., Mitoh, Y., Kohjitani, A., & Matsuo, R. (2003). Role of the hyperpolarization-activated cation current (Ih) in pacemaker activity in area postrema neurons of rat brain slices. The Journal of physiology, 552(1), 135–148.

29. Galvis, D., Wu, W., Hyson, R. L., Johnson, F., & Bertram, R. (2018). Interhemispheric dominance switching in a neural network model for birdsong. Journal of neurophysiology, 120(3), 1186–1197.

30. Gibb, L., Gentner, T. Q., & Abarbanel, H. D. (2009a). Brain stem feedback in a computational model of birdsong sequencing. Journal of neurophysiology, 102(3), 1763–1778.

31. Gibb, L., Gentner, T. Q., & Abarbanel, H. D. (2009b). Inhibition and recurrent excitation in a computational model of sparse bursting in song nucleus HVC. Journal of neurophysiology, 102(3), 1748–1762.

32. Glaze, C. M., & Troyer, T. W. (2013). Development of temporal structure in zebra finch song. Journal of neurophysiology, 109(4), 1025–1035.

33. Golowasch, J., Buchholtz, F., Epstein, I. R., & Marder, E. (1992). Contribution of individual ionic currents to activity of a model stomatogastric ganglion neuron. Journal of neurophysiology, 67(2), 341–349.

34. Golowasch, J., Casey, M., Abbott, L., & Marder, E. (1999). Network stability from activity-dependent regulation of neuronal conductances. Neural computation, 11(5), 1079–1096.

35. Golowasch, J., & Marder, E. (1992). Ionic currents of the lateral pyloric neuron of the stomatogastric ganglion of the crab. Journal of neurophysiology, 67(2), 318–331.

36. Gross, C., Yao, X., Engel, T., Tiwari, D., Xing, L., Rowley, S., Danielson, S. W., Thomas, K. T., Jimenez-Mateos, E. M., & Schroeder, L. M. (2016). MicroRNA-mediated downregulation of the potassium channel Kv4. 2 contributes to seizure onset. Cell reports, 17(1), 37–45.

37. Hahnloser, R. H., Kozhevnikov, A. A., & Fee, M. S. (2002). An ultra-sparse code underliesthe generation of neural sequences in a songbird. Nature, 419(6902), 65–70.

38. Hamaguchi, K., Tanaka, M., & Mooney, R. (2016). A distributed recurrent network contributes to temporally precise vocalizations. Neuron, 91(3), 680–693.

39. Harvey, C. D., Coen, P., & Tank, D. W. (2012). Choice-specific sequences in parietal cortex during a virtual-navigation decision task. Nature, 484(7392), 62–68.

40. Hodgkin, A. L., & Huxley, A. F. (1952). A quantitative description of membrane current and its application to conduction and excitation in nerve. The Journal of physiology, 117(4), 500.

41. Huguenard, J. (1996). Low-threshold calcium currents in central nervous system neurons. Annual review of physiology, 58(1), 329–348.

42. Jagodic, M. M., Pathirathna, S., Joksovic, P. M., Lee, W., Nelson, M. T., Naik, A. K., Su, P., Jevtovic-Todorovic, V., & Todorovic, S. M. (2008). Upregulation of the T-type calcium current in small rat sensory neurons after chronic constrictive injury of the sciatic nerve. Journal of neurophysiology, 99(6), 3151–3156.

43. Jin, D. Z. (2009). Generating variable birdsong syllable sequences with branching chain networks in avian premotor nucleus HVC. Physical Review E, 80(5), 051902.

44. Jin, D. Z., Ramazanoğlu, F. M., & Seung, H. S. (2007). Intrinsic bursting enhances the robustness of a neural network model of sequence generation by avian brain area HVC. Journal of computational neuroscience, 23, 283–299.

45. Kornfeld, J., Benezra, S. E., Narayanan, R. T., Svara, F., Egger, R., Oberlaender, M., Denk, W., & Long, M. A. (2017). EM connectomics reveals axonal target variation in a sequence-generating network. Elife, 6, e24364.

46. Kosche, G., Vallentin, D., & Long, M. A. (2015). Interplay of inhibition and excitation shapes a premotor neural sequence. Journal of Neuroscience, 35(3), 1217–1227.

47. Kozhevnikov, A. A., & Fee, M. S. (2007). Singing-related activity of identified HVC neurons in the zebra finch. Journal of neurophysiology, 97(6), 4271–4283.

48. Kubota, M., & Saito, N. (1991). Sodium-and calcium-dependent conductances of neurones in the zebra finch hyperstriatum ventrale pars caudale in vitro. The Journal of physiology, 440(1), 131–142.

49. Kubota, M., & Taniguchi, I. (1998). Electrophysiological characteristics of classes of neuron in the HVc of the zebra finch. Journal of neurophysiology, 80(2), 914–923.

50. Lee, A. K., & Wilson, M. A. (2002). Memory of sequential experience in the hippocampus during slow wave sleep. Neuron, 36(6), 1183–1194.

51. Leonardo, A., & Fee, M. S. (2005). Ensemble coding of vocal control in birdsong. Journal of Neuroscience, 25(3), 652–661.

52. Lewicki, M. S. (1996). Intracellular characterization of song-specific neurons in the zebra finch auditory forebrain. Journal of Neuroscience, 16(18), 5854–5863.

53. Li, M., & Greenside, H. (2006). Stable propagation of a burst through a one-dimensional homogeneous excitatory chain model of songbird nucleus HVC. Physical Review E, 74(1), 011918.

54. Llinás, R., & Yarom, Y. (1981). Properties and distribution of ionic conductances generating electroresponsiveness of mammalian inferior olivary neurones in vitro. The Journal of physiology, 315(1), 569–584.

55. Long, M. A., & Fee, M. S. (2008). Using temperature to analyse temporal dynamics in the songbird motor pathway. Nature, 456(7219), 189–194.

56. Long, M. A., Jin, D. Z., & Fee, M. S. (2010). Support for a synaptic chain model of neuronal sequence generation. Nature, 468(7322), 394–399.

57. MacDonald, C. J., Lepage, K. Q., Eden, U. T., & Eichenbaum, H. (2011). Hippocampal “time cells” bridge the gap in memory for discontiguous events. Neuron, 71(4), 737–749.

58. Marder, E., & Goaillard, J.-M. (2006). Variability, compensation and homeostasis in neuron and network function. Nature Reviews Neuroscience, 7(7), 563–574.

59. Markowitz, J. E., Liberti III, W. A., Guitchounts, G., Velho, T., Lois, C., & Gardner, T. J. (2015). Mesoscopic patterns of neural activity support songbird cortical sequences. PLoS biology, 13(6), e1002158.

60. Mooney, R. (2000). Different subthreshold mechanisms underlie song selectivity in identified HVc neurons of the zebra finch. Journal of Neuroscience, 20(14), 5420–5436.

61. Mooney, R. (2009). Neurobiology of song learning. Current opinion in neurobiology, 19(6), 654–660.

62. Mooney, R., Hoese, W., & Nowicki, S. (2001). Auditory representation of the vocal repertoire in a songbird with multiple song types. Proceedings of the national academy of sciences, 98(22), 12778–12783.

63. Mooney, R., & Prather, J. F. (2005). The HVC microcircuit: the synaptic basis for interactions between song motor and vocal plasticity pathways. Journal of Neuroscience, 25(8), 1952–1964.

64. Nottebohm, F., Paton, J. A., & Kelley, D. B. (1982). Connections of vocal control nuclei in the canary telencephalon. Journal of Comparative Neurology, 207(4), 344–357.

65. Pape, H.-C. (1996). Queer current and pacemaker: the hyperpolarization-activated cation current in neurons. Annual review of physiology, 58(1), 299–327.

66. Pastalkova, E., Itskov, V., Amarasingham, A., & Buzsaki, G. (2008). Internally generated cell assembly sequences in the rat hippocampus. Science, 321(5894), 1322–1327.

67. Pedarzani, P., McCutcheon, J. E., Rogge, G., Jensen, B. S., Christophersen, P., Hougaard, C., Strøbæk, D., & Stocker, M. (2005). Specific enhancement of SK channel activity selectively potentiates the afterhyperpolarizing current IAHP and modulates the firing properties of hippocampal pyramidal neurons. Journal of Biological Chemistry, 280(50), 41404–41411.

68. Peters, A. J., Chen, S. X., & Komiyama, T. (2014). Emergence of reproducible spatiotemporal activity during motor learning. Nature, 510(7504), 263–267.

69. Prather, J. F., Peters, S., Nowicki, S., & Mooney, R. (2008). Precise auditory–vocal mirroring in neurons for learned vocal communication. Nature, 451(7176), 305–310.

70. Roberts, T. F., Hisey, E., Tanaka, M., Kearney, M. G., Chattree, G., Yang, C. F., Shah, N. M., & Mooney, R. (2017). Identification of a motor-to-auditory pathway important for vocal learning. Nature neuroscience, 20(7), 978–986.

71. Schmitt, L. I., Wimmer, R. D., Nakajima, M., Happ, M., Mofakham, S., & Halassa, M. M. (2017). Thalamic amplification of cortical connectivity sustains attentional control. Nature, 545(7653), 219–223.

72. Shea, S. D., Koch, H., Baleckaitis, D., Ramirez, J.-M., & Margoliash, D. (2010). Neuron-specific cholinergic modulation of a forebrain song control nucleus. Journal of neurophysiology, 103(2), 733–745.

73. Shmiel, T., Drori, R., Shmiel, O., Ben-Shaul, Y., Nadasdy, Z., Shemesh, M., Teicher, M., & Abeles, M. (2006). Temporally precise cortical firing patterns are associated with distinct action segments. Journal of neurophysiology, 96(5), 2645–2652.

74. Skaggs, W. E., McNaughton, B. L., Wilson, M. A., & Barnes, C. A. (1996). Theta phase precession in hippocampal neuronal populations and the compression of temporal sequences. Hippocampus, 6(2), 149–172.

75. Terman, D., Rubin, J. E., Yew, A., & Wilson, C. (2002). Activity patterns in a model for the subthalamopallidal network of the basal ganglia. Journal of Neuroscience, 22(7), 2963–2976.

76. Vallentin, D., Kosche, G., Lipkind, D., & Long, M. A. (2016). Inhibition protects acquired song segments during vocal learning in zebra finches. Science, 351(6270), 267–271.

77. Varela, J. A., Sen, K., Gibson, J., Fost, J., Abbott, L., & Nelson, S. B. (1997). A quantitative description of short-term plasticity at excitatory synapses in layer 2/3 of rat primary visual cortex. Journal of Neuroscience, 17(20), 7926–7940.

78. Wang, C. Z. H., Herbst, J. A., Keller, G. B., & Hahnloser, R. H. R. (2008). Rapid interhemispheric switching during vocal production in a songbird. PLoS biology, 6(10), e250.

79. Wang, P., Liu, H., Wang, L., & Gao, R. X. (2018). Deep learning-based human motion recognition for predictive context-aware human-robot collaboration. CIRP annals, 67(1), 17–20.

80. Wang, X.-J., Liu, Y., Sanchez-Vives, M. V., & McCormick, D. A. (2003). Adaptation and temporal decorrelation by single neurons in the primary visual cortex. Journal of neurophysiology, 89(6), 3279–3293.

81. Weber, A. P., & Hahnloser, R. H. R. (2007). Spike correlations in a songbird agree with a simple Markov population model. PLoS computational biology, 3(12), e249.

82. Wild, J. M., Williams, M. N., Howie, G. J., & Mooney, R. (2005). Calcium-binding proteins define interneurons in HVC of the zebra finch (Taeniopygia guttata). Journal of Comparative Neurology, 483(1), 76–90.

83. Williams, A. H., O’Leary, T., & Marder, E. (2013). Homeostatic regulation of neuronal excitability. Scholarpedia, 8(1), 1656.

84. Yao, H., Donnelly, D. F., Ma, C., & LaMotte, R. H. (2003). Upregulation of the hyperpolarization-activated cation current after chronic compression of the dorsal root ganglion. Journal of Neuroscience, 23(6), 2069–2074.

85. Yu, A. C., & Margoliash, D. (1996). Temporal hierarchical control of singing in birds. Science, 273(5283), 1871–1875.

